# Heat-killed probiotic *Levilactobacillus brevis* MKAK9 and its exopolysaccharide promote longevity by modulating aging hallmarks and enhancing immune responses in *Caenorhabditis elegans*

**DOI:** 10.1101/2024.07.09.602715

**Authors:** Arun Kumar, Manti Kumar Saha, Vipin Kumar, Anupam Bhattacharya, Sagar Barge, Ashis K. Mukherjee, Mohan C. Kalita, Mojibur R. Khan

## Abstract

**Background:** Proteostasis is a critical aging hallmark responsible for removing damaged or misfolded proteins and their aggregates by improving proteasomal degradation through the autophagy-lysosome pathway (ALP) and the ubiquitin-proteasome system (UPS). Research on the impact of heat-killed probiotic bacteria and their structural components on aging hallmarks and innate immune responses is scarce, yet enhancing these effects could potentially delay age-related diseases.

**Results:** This study introduces a novel heat-killed *Levilactobacillus brevis* strain MKAK9 (HK MKAK9), along with its exopolysaccharide (EPS), demonstrating their ability to extend longevity by improving proteostasis and immune responses in wild-type *Caenorhabditis elegans*. We elucidate the underlying mechanisms through a comprehensive approach involving mRNA- and small RNA sequencing, proteomic analysis, lifespan assays on loss-of-function mutants, and quantitative RT-PCR. Mechanistically, HK MKAK9 and its EPS resulted in downregulation of the insulin-like signaling pathway in a DAF-16-dependent manner, enhancing protein ubiquitination and subsequent proteasomal degradation through activation of the ALP pathway, which is partially mediated by microRNA mir-243. Importantly, autophagosomes engulf ubiquitinylated proteins, as evidenced by increased expression of the autophagy receptor *sqst-3*, and subsequently fuse with lysosomes, facilitated by increased levels of the lysosome-associated membrane protein (LAMP) lmp-1, suggesting the formation of autolysosomes for degradation of the selected cargo. Moreover, HK MKAK9 and its EPS activated the p38 MAPK pathway and its downstream SKN-1 transcription factor, which are known to regulate genes involved in innate immune response (*thn*-1, *ilys*-1, *cnc-2*, *spp-9*, *spp*-21, *clec*-47, and *clec*-266) and antioxidation (*sod-3* and *gst-44*), thereby reducing the accumulation of reactive oxygen species (ROS) at both cellular and mitochondrial levels. Notably, SOD-3 emerged as a transcriptional target of both DAF-16 and SKN-1 transcription factors.

**Conclusion:** Our research sets a benchmark for future investigations by demonstrating that heat-killed probiotic and its specific cellular component, EPS, can downregulate the insulin-signaling pathway, potentially improving the autophagy-lysosome pathway (ALP) for degrading ubiquitinylated proteins and promoting organismal longevity. Additionally, we discovered that increased expression of microRNA mir-243 regulates insulin-like signaling and its downstream ALP pathway. Our findings also indicate that postbiotic treatment may bolster antioxidative and innate immune responses, offering a promising avenue for interventions in aging-related diseases.

## 1. Introduction

Advancements in medical science have significantly increased life expectancy worldwide, leading to a demographic shift characterized by a growing aging population. However, this demographic transformation brings with it a substantial economic burden, as the prevalence of age-associated diseases necessitates heightened healthcare investment [1]. Moreover, aging often correlates with a decline in immune function, rendering the elderly more susceptible to infections and other immune-related conditions [1]. According to projections by the United Nations, by 2050, one in every six individuals will be aged over 65, representing an estimated 1.5 billion people compared to 703 million in 2019 [2]. Consequently, there is a pressing need for research focused on understanding longevity, bolstering immune responses, and delaying the onset of age-related ailments, attracting global attention to this critical issue.

Previous research efforts have identified 12 hallmarks of aging, categorizing them into primary (loss of proteostasis, disabled macroautophagy, epigenetic changes, telomere shortening, and genomic instability), antagonistic (mitochondrial dysfunction, impaired nutrient sensing, and increased cellular senescence), and integrative categories (compromised intercellular communication, chronic inflammation, dysbiosis, and exhaustion of stem cells) [3]. Primary hallmarks initiate the accumulation of damage, while antagonistic hallmarks exert dual effects whose intensity can exacerbate negative outcomes [4]. Integrative hallmarks emerge when tissue homeostasis mechanisms can no longer compensate for the accumulation of damage [4]. Collectively, these factors drive the age-related decline in organ functionality. Additionally, dysregulation of immune responses can further exacerbate age-related decline and contribute to the development of various age-associated diseases. Consequently, strategies for aging intervention should target these key hallmarks, potentially delaying or mitigating the onset of age-related diseases as well as decline in age-associated immune responses. It is crucial to address both aspects of aging to enhance overall health and well-being in the elderly population.

Focusing on proteostasis, it is essential for cellular function, involving the balance between protein synthesis, folding, and degradation [5]. The proteostasis network, comprising proteasome degradation, autophagy, and chaperone-mediated folding, plays a critical role in maintaining protein homeostasis or known as proteostasis [6]. Firstly, proteasomal degradation mainly involves two major pathways for eliminating the damaged or misfolded proteins such as the ubiquitin-proteasome system (UPS) and the autophagy-lysosomal pathway (ALP) [7]. Notably, the proteasome activation is higher in centenarians and also delays aging in *in vivo* and *in intro* in model organisms [8]. In brief, protein ubiquitination tags regulatory and structural proteins by adding ubiquitin (Ub) (poly-Ub, 76 amino acid polypeptides covalently attached to target proteins) through the actions of three classes of enzymes (E1, E2, and E3) [9,

10]. The ubiquitinylated proteins are further degraded by 26S proteosome in UPS and autolysosomes (fusion of autophagosomes and lysosomes) in ALP [9, 10]. Dysfunction in the UPS or ALP pathways contributes to age-related diseases, highlighting their importance in healthy aging [9]. Secondly, the upregulation of autophagy is sufficient for promoting longevity in worms, flies, and rodents [11]. Autophagy is involved in lysosome-mediated degradation of damaged organelles and misfolded proteins [12]. Autophagosomes can engulf misfolded protein aggregates and damaged organelles and fuse them with lysosomes to form autolysosomes, which mediate degradation of the selected cargo [18]. Lysosome-associated membrane protein (LAMP) plays a crucial role in aiding the movement of autophagocytosed ubiquitinated proteins into the lysosome lumen for degradation [13]. Previous report suggested that the expression of LAMP decreases with aging and increasing transgenic expression can slow the aging process of the liver in mice [14]. Hence, the autophagy-lysosome pathway emerges as a vital strategy for eliminating intracellular damaged proteins and their aggregates. Lastly, epigenetic modifications, including non-coding RNA (miRNAs) molecules contribute to the aging process by regulating gene expression and affect the cellular mechanisms involved in longevity [15].

The gut microbiome’s dysbiosis, another aging hallmark, plays a vital role in aging across species, with nutritional strategies such as probiotic supplementation showing promise in promoting longevity [16]. Probiotics, comprising live microorganisms that confer health benefits when consumed in adequate quantities, are primarily represented by lactic acid bacteria (LAB), notably *Levilactobacillus* and *Bifidobacterium* [17]. Their mechanisms of action encompass positive modulation of gut microbiota, antimicrobial effects, enhancement of immune function, mucosal integrity improvement, anti-inflammatory properties, and immunomodulation [18]. Yet, the connections between probiotics, microRNA (miRNA) expression, proteostasis, and longevity need further exploration to fully understand their integrated impact on aging pathways and functionality of immune responses.

Studies exploring dietary interventions’ anti-aging effects have utilized diverse invertebrate to vertebrate model organisms, revealing conserved longevity mechanisms across species [19–21]. Notably, the nematode *Caenorhabditis elegans*, with its established symbiotic relationship with microbes and shared longevity mechanisms with humans, has been instrumental in aging research [22]. The major mechanisms include the p38 mitogen-activated protein kinase (p38 MAPK) pathway, the insulin-like signaling (IIS) pathway (DAF-2/DAF- 16), and transforming growth factor-β (TGF-β) signaling [23–26]. Research on *C. elegans* has also contributed insights into the role of immune responses in the aging process [23–26]. This understanding of how these pathways interact with immune function offers valuable insights into the comprehensive mechanisms underlying aging and identifies potential targets for anti-aging interventions.

While previous studies have primarily focused on live probiotic bacteria, exploring the role of postbiotics, such as components derived from heat-killed probiotics could provide valuable insights into their modulation of immune responses and aging-associated molecular pathways. In this context, we present findings demonstrating that a heat-killed probiotic *Levilactobacillus brevis* strain MKAK9 (HK MKAK9) and its exopolysaccharide (EPS) downregulate the insulin-like signaling pathway and enhancing proteasomal degradation (proteostasis) through ALP pathway, thereby promoting longevity in *C. elegans*. Additionally, the HK MKAK9-mediated activation of ALP is partially regulated by microRNA mir-243. Additionally, the treatment upregulates the p38 MAPK pathway to enhance immune responses and antioxidative machinery, thereby improving healthy aging in worms. This communication underscores the potential of HK MKAK9 and its EPS to positively influence diverse aspects of healthy aging, positioning them as promising postbiotic candidates for mitigating age-associated diseases.

## 2. Material and Methods

### 2.1 Materials

The bacterial growth media, including Luria-Bertini (LB) agar, deMan, Rogosa and Sharpe (MRS), and Nutrient agar, were purchased from Himedia, India. The bacterial standard food *Escherichia coli* OP50 (referred as OP50), hermaphrodite wild-type *C. elegans* strain (N2) and its mutants, including EU1 *skn-1* (zu67), KU25 *pmk-1* (km25), AU1 *sek-1* (ag1, AU3 *nsy-1* (ag3), GR1307 *daf-16* (mgDf50), NU3 *dbl-1*(nk3) V, DR1572 *daf-2* (e1368), RB2375 *lmp-1* (ok3228) X, CGC147 mir-1818, MT15021 mir-78(n4637) IV, MT15454 mir-243(n4759) IV, and MT16060 nDf64 V (mir-253), were obtained from Caenorhabditis Genetics Center (CGC), University of Minnesota, USA. The RNA from *C. elegans* was isolated using an RNA extraction kit (RNAeasy Mini Kit, Invitrogen, USA). Further, complementary DNA (cDNA) was prepared using a cDNA synthesis kit (Verso, Thermo Scientific, USA).

The human colon epithelial cell line HT-29 was obtained from the National Center of Cell Sciences (NCCS), Pune, India. The pathogenic strains, including *Staphylococcus aureus* MTCC 3160 and *E. coli* MTCC 1687, were obtained from the Microbial Type Culture Collection (MTCC), CSIR-Institute of Microbial Technology (CSIR-IMTECH), Chandigarh, India. The genomic extraction kit for isolating bacterial DNA was purchased from Sigma, Germany.

### 2.2. Isolation and bacterial growth conditions

The isolate MKAK9 was isolated from a mixed pickle sample collected from a local market in Maligaon, Guwahati, India. The strain MKAK9 was cultured on MRS agar at 30 °C for 48 h. A single streaked colony of strain MKAK9 was inoculated in MRS broth medium and incubated at 30 °C for 24 h, while OP50 was cultured in the LB broth at 37 °C for 12 h. The pathogens *S. aureus* MTCC 3160 and *E. coli* MTCC 1687 were obtained from the Microbial Type Culture Collection (MTCC), CSIR-Institute of Microbial Technology, Chandigarh, India. These strains were grown in a Nutrient broth medium at 37 °C for 12 h. All the inoculated bacterial strains were grown at 150 rpm in a shaking incubator (Orbital Shaker, India).

### 2.3 Characterization of the bacterium

The bacterial genomic extraction kit (Sigma, Darmstadt, Germany) was used to extract the genomic DNA of MKAK9, and genomic DNA was subjected to a polymerase chain reaction (PCR) to amplify full-length 16S ribosomal DNA using the eubacteria-specific universal primer 27F/1492R pair [27]. A total reaction mixture was prepared, and the PCR reaction was performed as described by Kumar et al. [27]. The sequencing of purified 16S rDNA fragments was outsourced to Macrogen Inc. (South Korea). The 16S rDNA sequence of MKAK9 was aligned with the sequences of the type strain *L. brevis* strain ATCC 14869 (Accession No. NR 044704.2) (Supplementary file 1: Table S1) and other strains’ available sequences (Supplementary file 1: Figure S1) within the National Center for Biotechnology Information (NCBI) database (Bethesda, USA).

### 2.4 Phylogenetic tree

The 16S rDNA sequence of the strain MKAK9 was compared with type strain *Levilactobacillus brevis* strain ATCC 14869 (Accession no.: NR 044704), other related *Lactobacillus* species, and with an outgroup genera *E. faecium* ATCC 19434. The phylogenetic tree was reconstructed using the neighbour-joining algorithms. The phylogenetic neighbours were identified, and similarity was calculated to align them using phylogenetic tree analysis (Mega 7 version). The bootstrap analysis measured the branching’s confidence intervals and branch-length errors. The nodes consist of bootstrap values (percentages of 1000 replications) greater than 60%.

### 2.5 Screening for probiotic properties

#### 2.5.1 Probiotic screening tests

The probiotic screening tests for the strain MKAK9, including survivability to gastrointestinal transit, bile salt tolerance, and adhesion to intestinal epithelial cell line (HT-29), were performed using protocols previously described by Kumar et al. [27]. Each experiment was conducted thrice, and each time performed with three replicates.

#### 2.5.2 Antibiotic sensitivity assay

A Hexa G-PLUS antibiotic ring (Himedia, India) was used to perform an antibiotic sensitivity assay. The antibiotic ring includes tetracycline, ampicillin (10 μg), sulphatriad (300 μg), chloramphenicol (25 μg), streptomycin (10 μg), tetracycline (25 μg), and penicillin-G (10 μg). The grown MKAK9 culture was briefly adjusted to 10^9^ CFU/ml and spread-plated on MRS agar plates. The antibiotic ring was placed aseptically on the plate, and their zone of inhibition was measured after 48 h.

### 2.6 Lifespan assay

The Nematode Growth Medium (NGM) plates with sufficient eggs of worms were synchronized using a protocol described by Stiernagle [28]. The synchronized eggs were allowed to hatch and grown on standardized 10^9^ CFU/mL concentration of *E. coli* OP50-seeded NGM plates at 20 °C for 48 h. The lifespan assays were performed on young adult (L4 worms) wild-type N2 and mutant worms. The bacterial culture with standardized 10^9^ CFU/mL bacterial cells in M9 buffers was heated for 2 h at 70 °C. The lifespan experiment was performed with three technical replicates in which 50 worms were transferred individually to previously bacterial-seeded NGM plates and incubated at 20 °C. The number of worms per plate was recorded every 24 h and transferred once every two days to new NGM plates seeded with heat-killed bacterial food. The survival analysis was conducted in OASIS 2 software using the Kaplan-Meier method, and the mean lifespan was calculated.

### 2.7 Developmental rate assay

The effect of heat-killed bacterial diets on the developmental rate of worms was analyzed using a protocol discussed by Soukas et al. [29]. In brief, age-synchronized 20 young adult worms were transferred to both bacterial-seeded NGM plates and observed every 30 min until the first laid eggs under a stereo zoom microscope (SMZ1270, Nikon, Japan). The experiment was performed in triplicate, and the experiment was conducted twice.

### 2.8 Measurement of body length

The body length of wild-type N2 worms on heat-killed OP50 and MKAK9 was measured by a procedure described by Kumar et al. [27]. Ten worms were imaged under a stereo zoom microscope (SMZ1270, Nikon, Japan) for each heat-killed bacterial strain, and the experiment was performed in triplicate for each heat-killed bacterium.

### 2.9 Determination of locomotory activity and pharynx pumping

The locomotory activity and pharynx pumping of worms were measured using a protocol described by Kumar et al. [27]. The synchronized young adult worms were cultured on NGM plates seeded with heat-killed OP50 or MKAK9 for 14 days. The locomotory activity was determined by counting the number of body bendings of each worm per minute. The pharynx pumps were measured by counting the number of pharynx pumps for 30 seconds per individual worm. The locomotory activity and pharynx pumping of heat-killed OP50 or MKAK9-treated worms were recorded on day 14 using a stereo zoom microscope (S8 APO E, Leica, Germany).

Ten worms were accessed for each heat-killed bacterial strain, and the experiment was performed in triplicates for each heat-killed bacterium.

### 2.10 Accumulation of the aging pigment lipofuscin

The accumulation of the aging pigment lipofuscin was determined in worms according to the protocol described by Kumar et al. [27]. Ten worms were imaged under a confocal microscope (TCS SPE, Leica, Germany) for each heat-killed bacterial strain, and the experiment was performed in triplicate for each heat-killed bacterium.

### 2.11 Resistance against pathogenic bacterial infections

The age-synchronized worms were pre-cultured on OP50 for 2 days, transferred to heat-killed OP50 or MKAK9 for 24 h and maintained at 20°C. Then, worms were individually transferred to each NGM plate seeded with Gram-positive pathogen *S. aureus* strain MTCC3160 (10^9^ CFU/mL) and Gram-negative pathogen *E. coli* strain MTCC 1687 (10^9^ CFU/mL). The experiment was performed with three technical replicates in which 50 worms and number of worms per plate were counted every 24 h and transferred once every two days to new NGM plates seeded with pathogens. The survival analysis was conducted in OASIS 2 software using the Kaplan-Meier method, and the mean survival days were calculated.

### 2.12 Determination of reduction in the colonization of pathogens in worm’s intestine

The age-synchronized young adult worms were cultured on heat-killed OP50 or MKAK9 for 24 h at 20 °C. The worms were then transferred to NGM plates seeded with the pathogens *S. aureus* strain 3160 and *E. coli* strain 1687. Ten worms per bacterial treatment over several days (1, 3, and 5 days) were collected, washed three times with M9 buffer, and treated with drops of gentamycin (25 μg/ml) to remove the remaining attached pathogen on the worm’s surface. Further, the worms were washed thrice with M9 buffer and mechanically lysed using a pestle in a 1.5 ml centrifuge tube. Then, the whole worm lysate was plated on their respective culture media plates. Each bacterial pathogen’s colony forming units (CFU) were counted after incubating at 37 °C for 24 h, and the number of colonized bacterial pathogens per worm was calculated. The experiment was performed in triplicate for each heat-killed bacterium.

### 2.13 Oxidative stress and thermotolerance assay

The age-synchronized young adult worms were individually transferred to heat-killed OP50 or MKAK9-seeded NGM plates and allowed to culture for 3 days at 20 °C. For the oxidative stress resistance assay, the pre-cultured worms on heat-killed OP50 or MKAK9 were individually transferred onto NGM plates supplemented with 10 mM paraquat. The pre-cultured worms on OP50 or MKAK9 were incubated at 35 °C for the thermotolerance assay. The viable worms were counted until all worms were dead for both assays. The experiment was performed with three replicates for each heat-killed bacterium, and 50 worms were transferred to three plates per treatment.

### 2.14 Quantitative Reverse Transcription-Polymerase Chain Reaction (qRT-PCR) to measure the differential expression of selected genes in HK MKAK9-treated worms in comparison to HK OP50-treated worms

The age-synchronized worms were grown on OP50-seeded NGM plates for 2 days at 20 °C. These worms were transferred individually to the NGM plates seeded with HK OP50 or HK MKAK9 and maintained at 20 °C for 24 h. Approximately 500 worms per treatment group were collected and washed five times with M9 buffer to remove the remaining bacteria. The total RNA was extracted using the RNAeasy mini kit (Invitrogen, USA) and reverse transcribed to cDNA using the Verso cDNA synthesis Kit (Thermo Scientific, USA) as per the protocol described by Kumar et al. [27]. The primer sequences of the studied genes are listed in Table S2).

The qRT-PCR was performed to measure the differential expression of genes using SYBR Green (Applied Biosystems, USA) in a real-rime PCR machine (Bio-Rad Laboratories, Inc., CA, USA) [27]. In a 96-well plate, each well contained a total volume of 20 μl, including 1.2 μl of 10-fold diluted cDNA, 0.4 μl of each primer (10 μM), 10 μl of 2X SYBR Green, and 8 μl of deionized water (Millipore, MA, USA). The reaction conditions were setup in qRT-PCR as described by Kumar et al. [27], followed by a melt curve analysis. The experiment was performed independently twice with three replicates each time, and the relative expression of each candidate gene was analyzed using the change in 2^-ΔΔCt^ method. *act-*1, a housekeeping gene, was used to normalize the expression of each gene in this study. The change in comparative Ct (Ct = Threshold cycle value) for each gene in the sample was calculated using ΔCtSample = CtSample – Ctact-1. The final change in Ct value between the test and control sample was calculated as 2^-ΔΔCt^, where ΔΔCt = ΔCtHK MKAK9 – ΔCtHK OP50.

### 2.15 mRNA and small RNA sequencing analysis

The young adult age-synchronized wild-type N2 worms were treated with heat-killed OP50 or MKAK9 at 20 °C for 24 h. For mRNA and small RNA sequencing, the total RNA was extracted from three biological replicates per group using the PureLink^TM^ RNA Mini-Kit as per the manufacturer’s instructions. The concentration and quality of RNA were assessed using a Qubit fluorometer. The samples containing an RNA integrity number (RIN) ≥ 7 were checked using the Agilent Bioanalyzer 2100 (Agilent Technologies, USA) and considered qualified for library preparation. mRNA and small RNA sequencing was performed on the Illumina Hiseq 2500 and Illumina NextSeq 500 platforms, respectively, with commercial sequencing service provider MedGenome Labs Ltd., Bengaluru, Karnataka, India. The transcript levels were measured using fragments per kilobases per million reads and compared to different genes in the samples. The changes in the expression of genes were identified after filtering the mRNA sequencing data and microRNA sequencing with the cut-off values involving a two-fold change in the expression level between samples and a false discovery rate (FDR) analogue with a *p*-value less than 0.05.

The mRNA sequencing data gene list was then subjected to Gene Ontology analysis in the Kyoto Encyclopedia of Genes and Genomes (KEGG) and protein domain enrichment analysis. For gene ontology analysis, we have used “TopGO” and the genocide annotation package of *C. elegans* (*“*org.Ce.eg.db”, bioconductor) in R. We defined the significantly enriched terms annotated for each of the categories of BP (Biological process), MF (Molecular function), and CC (Cellular components) with a *p-*value < 0.05 for the Fisher exact test. We have used the ggplot2 package to plot and visualize the significant GO terms. KEGG and protein enrichment analysis were performed using DAVID Bioinformatics Resources 6.8. The enrichment score is defined as –log (*p*-value), and the terms with a *p*-value less than 0.05 were plotted. The hierarchical clustering was analyzed using heatmap.2 in the R package (v3.6.3).

### 2.16 Quantitative proteomics analysis of worms treated with heat-killed probiotic strains MKAK9 or OP50

The age-synchronized young adult wild-type N2 worms were transferred to NGM plates seeded with heat-killed OP50 or MKAK9 for 24 h and maintained at 20 °C. The treated worms were washed with M9 buffer to remove the residual bacteria, centrifuged, and resuspended in 50 mM Tris-HCl buffer containing a protease inhibitor cocktail. The worms were then sonicated on ice, and debris was removed by an additional step of centrifugation at 7,500 x g for 5 min, and the resulting supernatant was separated. The protein concentration was determined using extracted protein using Bradford reagent, and the protein concentration was maintained at 100 μg per sample. Briefly, 20 µg of the protein was reduced with dithiothreitol (DTT) and alkylated using 100 mM iodoacetamide (IAA), followed by precipitation with chilled acetone, and incubated for 4 h at 80 °C.

The precipitated protein was then digested with trypsin overnight at 37 °C. Formic acid (1%) was added to stop the reaction, and samples were dried and desalted to remove buffer salts and detergents. The digested peptides were then subjected to a nano-Orbitrap Fusion Tribrid Mass Spectrometer coupled to a Thermo EASY nanoLC 1200 chromatographic system. MS and MS/MS scan ranges at 375 to 1700 m/z were performed. The raw data obtained from the MS/MS analysis was analyzed using MaxQuant, and parameters for identifying proteins were adopted from Bianchi et al. [30]. The spectra were searched in the UniProt reference proteome database against *C. elegans* (Taxonomy ID: 6239) and included a set of common contaminants. The matches for peptide spectra were adjusted to FDR of 0.01 by the target-decoy approach, respectively. Carbamidomethylation of cysteine was considered a static modification, and oxidation of protein N-terminal acetylation and methionine residues was included as a variable modification. The relative abundance of identified proteins in treated groups was calculated from spectral intensity, and fold change was calculated from the relative abundance of identified proteins.

### 2.17 Extraction of the cell wall and cytoplasmic fractions of strain MKAK9

The cell wall and cytoplasmic fractions were prepared and discussed as described by Wang et al. [31]. The cytoplasmic fraction was used to extract both peptidoglycan-wall teichoic acid (PGN-WTA) and lipoteichoic acid (LTA) of the strain MKAK9 using a procedure described by Wang et al. [31]. The pellet was used to extract the S-layer protein of the strain MKAK9, which was processed as per the protocol of Eslami et al. [32]. On the other hand, the grown MKAK9 culture broth was boiled at 100 °C for 30 min to disperse the cell-bound EPS and neutralize endogenous enzymes. The broth culture was centrifuged (5000 g for 20 min at 4 °C), and the pellet and solidified proteins were separated. The resultant cell-free supernatant was used for extracting the EPS using a protocol described by Rajoka et al. [33].

To perform the longevity assay with cell wall and cytoplasmic fractions of strain MKAK9, the age-synchronized young adult wild-type worms were treated with 1% (v/v) of cell wall, cytoplasmic fraction, PGN-WTA, LTA, S-layer, or EPS fractions derived from strain MKAK9 bacterial cells as described by Wang et al. [31]. The remaining procedure for conducting the lifespan assay was the same as the protocol mentioned above in Section 2.6. For qRT-PCR, the age-synchronized young adult worms were treated with cell wall and cytoplasmic fractions for 24 h, and RNA was isolated for qRT-PCR.

### 2.18 GSH/GSSG Assay

The oxidized glutathione (GSSG) and reduced glutathione (GSH) were individually measured by employing the Glutathione Assay Kit (Promega, USA) as described by Nakagawa et al. [23].

### 2.19 SOD activity assays

SOD Assay Kit-WST (Sigma, USA) was employed to measure superoxide dismutase (SOD) activity in day-14 bacterial-treated worms, as described by Nakagawa et al. [23].

### 2.20 Measurement of Intracellular ROS

The intracellular ROS level was measured by employing 2′,7′-dichlorofluorescein diacetate (H2DCFDA) fluorescent dye in day-14 worms cultured on heat-killed OP50 or MKAK9, as per the protocol discussed by Kumar et al. [27].

### 2.20 Mitochondrial ROS staining

The mitochondrial ROS level was measured using MitoTracker® Red CM-H2XRos (Invitrogen, USA) in day-14 worms cultured on heat-killed OP50 and MKAK9, as per the protocol described by Kumar et al. [27].

### 2.21 Measurement of Adenosine Triphosphate (ATP)

The ATP was measured using the Roche ATP Bioluminescent HSII kit (Merck, Germany) as described by Nakagawa et al. [14]. Briefly, the lyophilized ATP was dissolved to a final concentration of 1 μM and further dissolved to make an ATP standard curve with a serial dilution of 1 μM to 1 x 10^-5^ μM. The preparation of the protein extract of day-14 worms grown on heat-killed OP50 or MKAK9 and bioluminescence were measured for each treatment group using a multimode reader (GloMax Discoverer Promega, Madison, WI, USA). ATP concentration for each treatment group was measured using a log-log plot of the standard curve and expressed as nmol of ATP concentration per mg protein.

### 2.23 Quantification of protein

Protein concentration was measured using the Pierce™ BCA Protein Assay Kit (Thermo Fisher Scientific, USA). Bovine serum albumin (BSA) was used as a standard protein to assess protein concentration.

### 2.24 Statistical analysis

The experiments were performed with three independent replicates, and all experiments were repeated twice. The data was depicted as the mean ± standard error mean (SEM). The statistical analysis was performed using the Online Application for Survival Analysis 2 (OASIS2) program for longevity and other survival assays. The log-rank test was used to evaluate the difference between the survival curves of worms. SigmaPlot version 12.0 (San Jose, CA, USA) performed all statistical computation analysis. The statistical analysis between treatment groups was performed using the Student’s t-test and a one-way analysis of variance (ANOVA). The mean ± SEM with a *p*-value <0.05 for all the experiments was considered statistically significant.

## 3. Results

### 3.1 Taxonomic characterization of the strain MKAK9 and its *in vitro* probiotic properties

The 16S rDNA sequence of the strain MKAK9 was sequenced, and subsequent BLAST analysis revealed significant similarity to the type strain *Levilactobacillus brevis* strain ATCC 14869 (Accession No. NR 044704.2). To establish phylogenetic relationships, a rooted phylogenetic tree of the strain MKAK9 was constructed alongside sequences of the type strain *L. brevis* ATCC 14869 and other *Levilactobacillus* species (Supplementary file 1, Fig. S1). The analysis indicated that the closest relative of strain MKAK9 was *Levilactobacillus brevis* ATCC 14869, exhibiting a remarkable 99% pairwise sequence similarity. Consequently, strain MKAK9 was officially submitted to the NCBI database as *Levilactobacillus brevis* strain MKAK9 (Supplementary file 1, Table S1).

Strain MKAK9 underwent rigorous *in vitro* screening tests for probiotic attributes, following the guidelines set forth by the Department of Biotechnology (DBT) and the Indian Council of Medical Research (ICMR), Government of India [34]. These *in vitro* tests encompassed evaluations for tolerance to acidic and bile salt conditions, as well as adherence to gastrointestinal epithelium. Notably, MKAK9 demonstrated robust viability, retaining 83.24% and 79.22% viability after 1 h of exposure to simulated gastric juice (SGJ) at pH 3.0 and pH 1.0, respectively (Supplementary file 1, Table S3). Furthermore, exposure to SGJ containing pepsin at pH 2.0 resulted in viability rates of 83.05% after 1 h and 79.08% after 3 h (Supplementary file 1, Table S3). Following treatment with SGJ containing pancreatin (pH 8.0) for 4 h, viability remained high at 81.86% (Supplementary file 1, Table S3). Bile salt tolerance tests demonstrated that strain MKAK9 exhibited remarkable resilience, retaining 85.19% and 79.89% survival rates against bile salt concentrations of 0.3% and 1%, respectively, after 4 h (Supplementary file 1, Table S3).

Given its promising tolerance to simulated gastrointestinal conditions, strain MKAK9 underwent further evaluation for adhesion using the gastrointestinal cell line HT-29. Strain MKAK9 exhibited a substantial adherence rate of 72.17% following a 4 h incubation period (Supplementary file 1, Table S3). Additionally, an antibiotic sensitivity assay revealed that strain MKAK9 was susceptible to tetracycline, ampicillin, sulphatriad, and streptomycin, while demonstrating resistance to penicillin-G and chloramphenicol (Supplementary file 1, Table S4). These findings align closely with the antibiotic sensitivity profiles reported for other *Levilactobacillus* strains, as described by Costa et al. [35].

### 3.2 Feeding of HK MKAK9 slowed the aging of worms

Treatment with live strain MKAK9 significantly increased the longevity of worms by 25.61% compared to the live standard bacterium *E. coli* OP50 (\*\*\**P <* 0.0001, log-rank test) (Fig. 1A). Notably, feeding heat-killed MKAK9 (abbreviated as HK MKAK9) also extended the mean lifespan of worms by 24.40% compared to HK OP50 (\*\*\**P <* 0.0001, log-rank test) (Fig. 1A). This suggests that both live and heat-killed MKAK9 can enhance the mean lifespan of worms, prompting us to proceed with HK MKAK9 for further investigation compared to HK OP50. Additionally, the longevity-promoting effect also supported the *in vivo* investigation in worms, suggesting that strain MKAK9 is beneficial and non-pathogenic to *in vivo* host.

**Figure 1.**
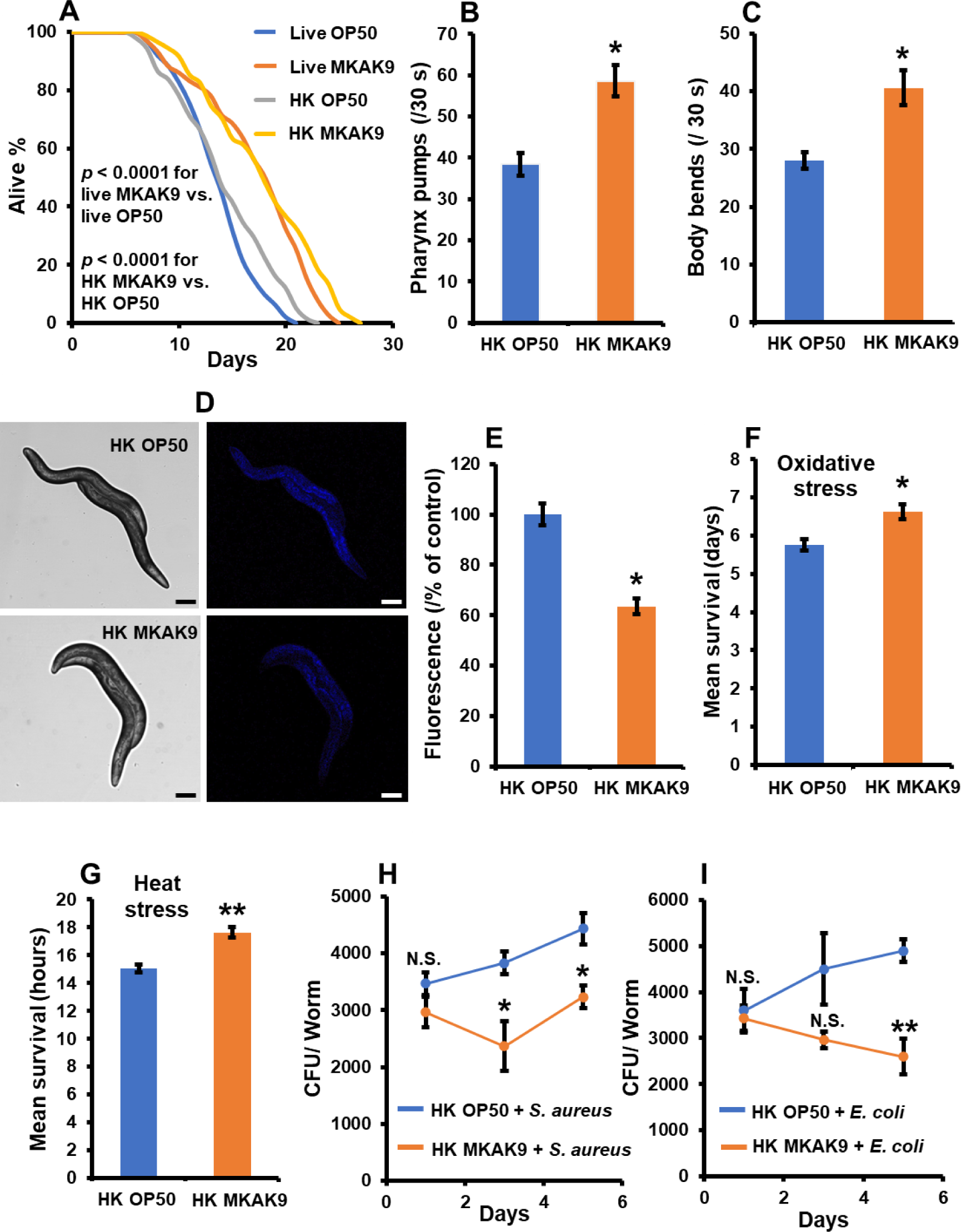
Feeding of probiotic heat-killed MKAK9 improves longevity, age-associated biomarkers and resistance of worms against pathogenic infections and abiotic stress in worms. (A) A lifespan assay on worms was performed after feeding with the bacteria: heat-killed *E. coli* OP50 (HK OP50), *E. coli* OP50 (OP50), heat-killed *L. brevis* MKAK9 (HK MKAK9), *L. brevis* MKAK9 (MKAK9), and (\*\*\**P <* 0.0001, log-rank test). (B-C) The pharynx pumps, and locomotory activity of heat-killed bacterium-treated worms were measured on day 14. (D-E) The accumulation of the aging pigment lipofuscin was measured on day 14 in heat-killed bacterium-treated worms. The lipofuscin levels were observed in worms under a confocal microscope (10X magnification, scale bar, 100 μm). (F-G) Treatment of HK MKAK9 increased the mean survival of worms against oxidative stress (Paraquat, 10 mM) and heat stress at 35 °C. (H-I) Pre-treatment of worms on HK reduced the colonization of pathogens *S. aureus* MTCC3160 and *E. coli* MTCC 1687 in the intestine of worms. The heat-killed bacterium-treated effects were statistically compared using a Student’s t-test (\**P <* 0.05 and \*\**P <* 0.01). The error bars represent the mean ± SEM.

Conducting a developmental rate assay on different bacterial diets revealed that HK MKAK9-fed worms exhibited slower development from eggs to the egg-laying reproductive adult stage compared to HK OP50-treated worms (\*\**P <* 0.01) (Supplementary file 1, Fig. S2). However, there were no significant changes observed in the body size of HK MKAK9-fed worms compared to HK OP50-fed worms (*P* > 0.05 on days 4, 5, 6, and 7) (Supplementary file 1, Fig. S3).

The longevity of organisms often correlates with changes in age-related biomarkers in worms, such as locomotory activity, pharyngeal pumping rate, and the accumulation of the aging pigment lipofuscin [22]. To explore the effect of HK MKAK9, we conducted analyses on these biomarkers. Firstly, we examined the effect of HK MKAK9 on the pharyngeal pumping rate. Our results revealed a notable enhancement in the pharyngeal pumping rate of worms by 52.55% compared to HK OP50-fed worms (\**P <* 0.05) (Fig. 1B). Subsequently, we evaluated the locomotory activity of worms treated with HK MKAK9 by quantifying the number of body bends per 30 seconds. Interestingly, we observed a substantial improvement in the frequency of body turns, which increased by 45.20% compared to HK OP50-fed worms (\**P <* 0.05) (Fig. 1C). Furthermore, we investigated the accumulation of the aging pigment lipofuscin in 14-day worms treated with HK MKAK9. Strikingly, our analysis unveiled a significant reduction of 36.5% in the lipofuscin levels in HK MKAK9-treated worms compared to HK OP50 (\**P <* 0.05) (Figs. 1D and E). These findings collectively suggest that HK MKAK9 slowed aging and exerted beneficial effects on age-related biomarkers, indicating its potential as a modulator of aging processes in worms.

### 3.3 Feeding of HK MKAK9 provided resistance against abiotic stress and pathogenic infections

The administration of HK MKAK9 significantly enhanced the survival of worms against oxidative stress (10 mM) by 15.10% compared to HK OP50-treated worms (\**P <* 0.05) (Fig. 1F). Furthermore, we investigated the protective effects of HK MKAK9 against thermal stress (35 °C) on worms. Our results revealed a notable 17.15% increase in the mean survival rate of HK MKAK9-treated worms compared to those fed with HK OP50 (\*\**P <* 0.01) (Fig. 1G).

We investigated the effect of HK MKAK9 on resistance against pathogenic infections in worms. Our findings revealed that HK MKAK9 significantly enhanced worm survival by 23.72% and 16.69% against *S. aureus* strain MTCC3160 and *E. coli* strain MTCC1687, respectively (\*\**P <* 0.001 for *S. aureus* and \**P <* 0.01 for pathogen *E. coli*, log-rank test) (Supplementary file 1, Figs. S4A and B). Furthermore, we explored whether HK MKAK9 could mitigate pathogenic bacterial colonization, thereby extending worm survival. To investigate this possibility, we pre-cultured young adult worms on either HK OP50 or HK MKAK9 and subsequently transferred them to pathogen-seeded NGM plates. Remarkably, our results demonstrated a significant reduction in the colonization of both *S. aureus* and *E. coli* in HK MKAK9-treated worms compared to those fed with HK OP50 (N.S. *P* > 0.05 for *S. aureus* on day 1, \**P <* 0.05 for *S. aureus* on day 3 and day 5, N.S. *P* > 0.05 for pathogen *E. coli* on day 1 and day 3, \*\**P <* 0.01 for pathogen *E. coli* on day 5) (Figs. 1H and I). Notably, the number of pathogenic bacteria exhibited a significant decrease after day 3 of pathogen treatment (\**P <* 0.05 for *S. aureus* on day 5 and \*\**P <* 0.01 for pathogen *E. coli* on day 5) (Figs. 1H and I).

### 3.4 Treatment of HK MRKA9 improved cellular and mitochondrial redox states in worms

We aimed to elucidate the alterations in cellular and mitochondrial redox status in HK MKAK9-treated worms. Firstly, employing 2,7-dichlorofluorescein diacetate (H2DCFDA), we assessed cytoplasmic ROS levels in vivo in HK MKAK9- and HK OP50-treated worms. Encouragingly, our results revealed a significant reduction of 45.2% in cytoplasmic ROS levels in HK MKAK9-treated worms compared to those treated with HK OP50 (\*\**P <* 0.01) (Figs. 2A and B). Subsequently, we evaluated superoxide dismutase (SOD) activity in worms following HK MKAK9 treatment. HK MKAK9 treatment led to a notable enhancement of 37.5% in SOD activity compared to HK OP50-treated worms (\**P <* 0.05) (Fig. 2C). Furthermore, we examined the glutathione (GSH) to glutathione disulfide (GSSG) ratio, a crucial indicator of cellular oxidative environment [36]. Intriguingly, HK MKAK9 treatment resulted in approximately a two-fold improvement in the GSH/GSSG ratio in day 14 worms compared to those treated with HK OP50 (\**P <* 0.05) (Fig. 2D).

**Figure 2.**
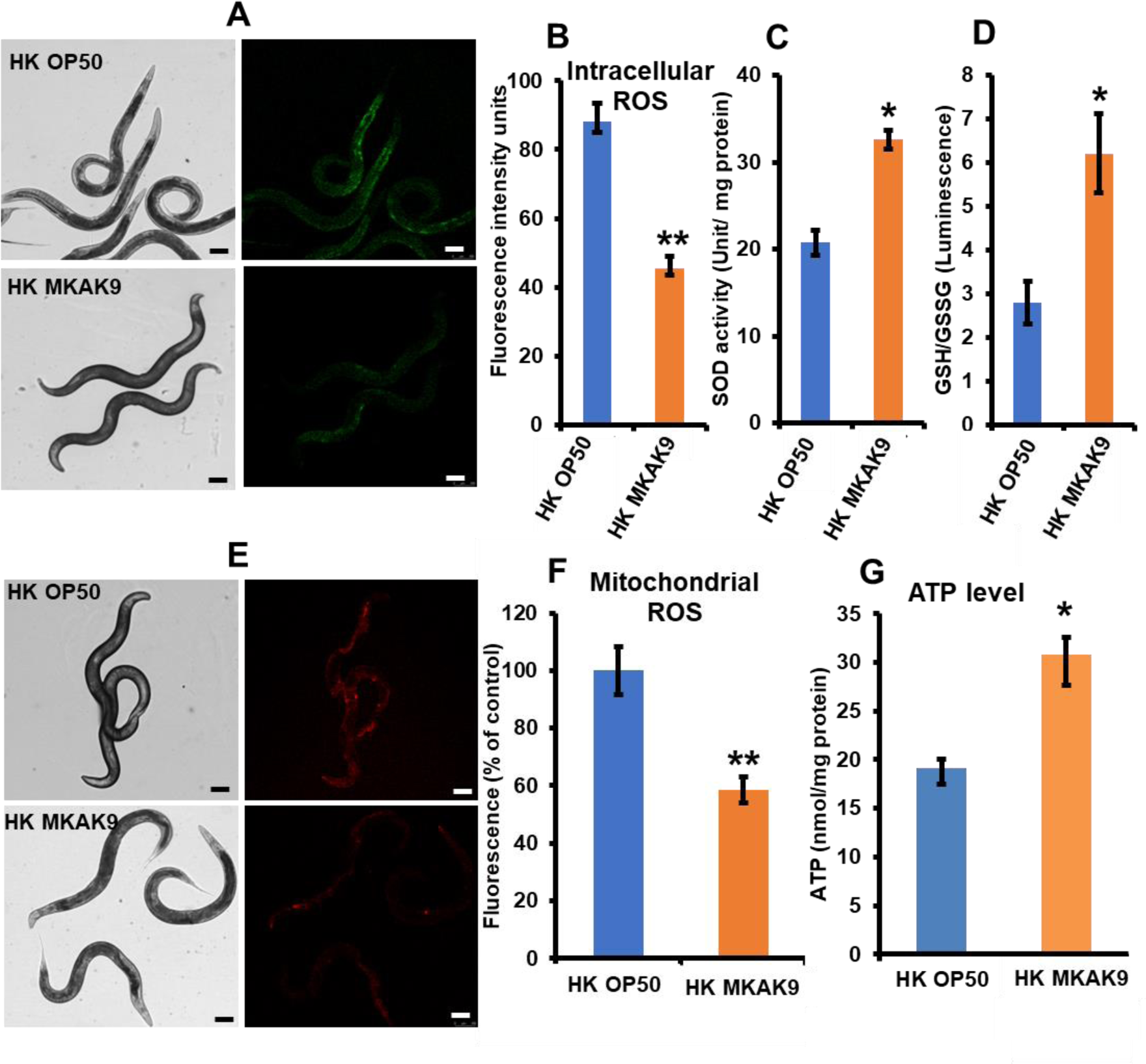
Treatment of HK MKAK9 improved the cellular and mitochondrial redox state in worms. (A-D) Feeding of HK MKAK9 reduced the accumulation of cytoplasmic ROS levels, improved SOD activity, and the GSH/GSSG ratio in worms. **(E-G)** Treatment of HK MKAK9 also reduced mitochondrial ROS levels, thereby improving ATP levels in worms compared to HK OP50-treated worms. The cytoplasmic and mitochondrial ROS levels were observed in worms under a confocal microscope (10X magnification, scale bar, 100 μm). The heat-killed bacterium-treated effects were statistically compared using Student’s t-test (N.S. for *P* > 0.05, \**P <* 0.05, and \*\**P <* 0.01). The bars represent the mean ± SEM.

We employed MitoTracker Red CMXRos to explore the changes in mitochondrial redox state. Our findings demonstrated a substantial reduction of 41.4% in mitochondrial ROS levels in HK MKAK9-treated worms compared to HK OP50-treated worms (\*\**P <* 0.01) (Figs. 2E and F). Moreover, we investigated the effect of HK MKAK9 on ATP synthesis levels in worms. HK MKAK9 treatment led to a significant improvement of 38.2% in ATP levels in day 14 worms compared to those treated with HK OP50 (\**P <* 0.05) (Fig. 2G). These results collectively showed the potent antioxidative effects of HK MKAK9 in mitigating oxidative stress and enhance mitochondrial function in worms.

### 3.5 Treatment of HK MKAK9 regulates evolutionary conserved insulin-like signaling and the p38 MAPK pathway to promote longevity and immune responses in worms

Diverse conserved signaling mechanisms govern aging. To elucidate the role of the p38 MAPK pathway in HK MKAK9-mediated longevity in worms, we examined its involvement using loss-of-function mutants of the p38 MAPK cascade (*nsy-1*, *sek-1*, and *pmk-1*). HK MKAK9 failed to enhance longevity in these mutants (*P* > 0.05 for *nsy-1*, *sek-1*, and *pmk-1*, log-rank test) (Table S5) (Figs 3A, B, and C). Additionally, we investigated whether HK MKAK9- induced upregulation of the p38 MAPK cascade activated downstream gene expression, specifically *skn-1*. Notably, the *skn-1* (zu67) allele mutant failed to exhibit HK MKAK9- induced longevity (*P* > 0.05, log-rank test) (Table S5) (Fig. 3D). Next, explored the involvement of insulin-like signaling, focusing on *daf-2* and *daf-16* genes, in mediating HK MKAK9-induced longevity. Both loss-of-function *daf-2* and *daf-16* mutants failed to extend longevity upon HK MKAK9 treatment (*P* > 0.05 for *daf-2* and *daf-16*, log-rank test) (Table S5) (Figs. 3E and F). Furthermore, we assessed the role of the TGF-β homolog *dbl-1*, finding that HK MKAK9 significantly extended longevity in the *dbl-1* mutant (\*\*\**P <* 0.0001, log-rank test) (Table S5) (Fig. 3G).

**Figure 3.**
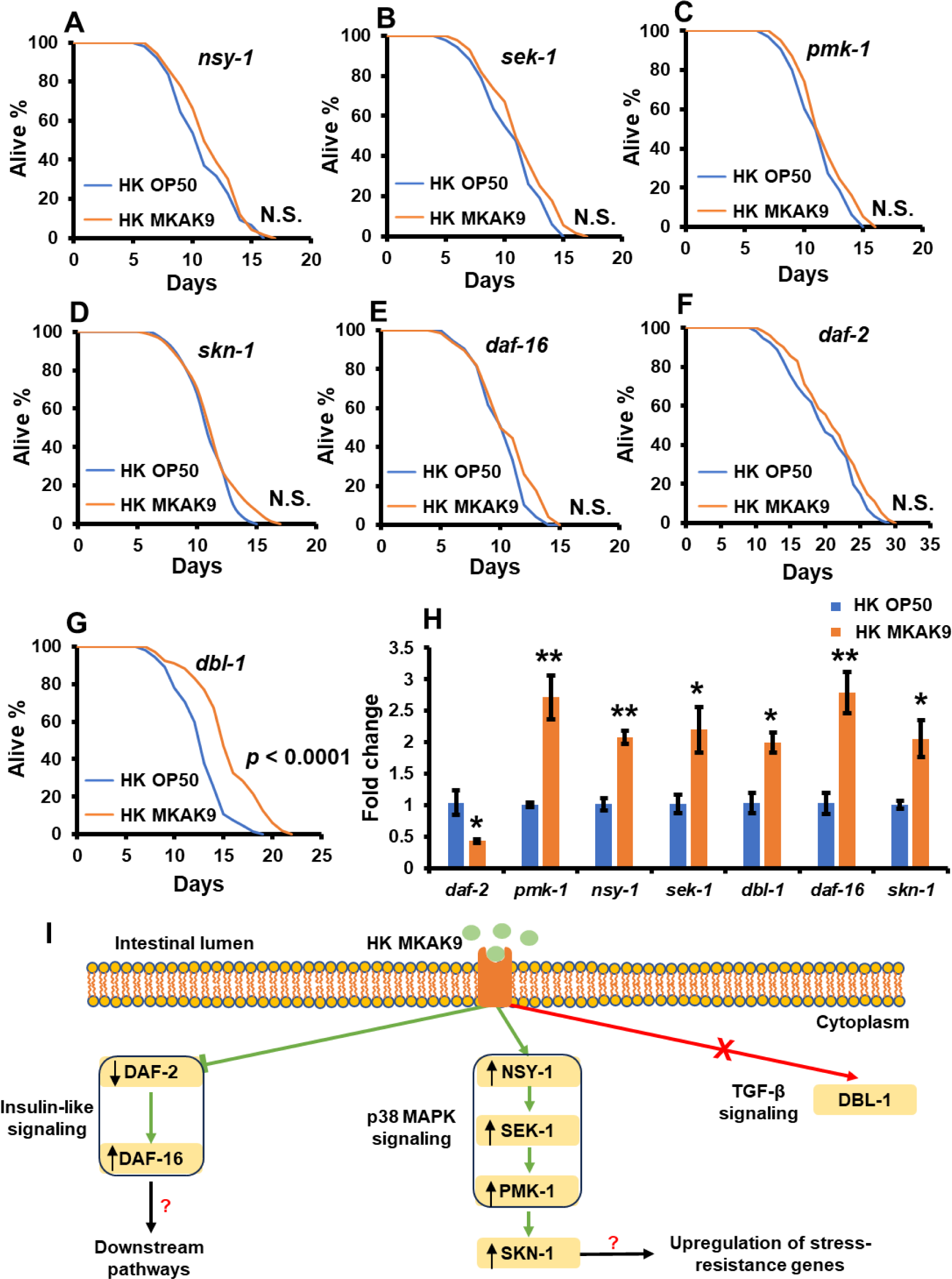
Effect of HK MKAK9 on the lifespan of *C. elegans* loss-of-function mutants. Longevity assays were conducted with mutants (**A**) *nsy-1* (ag3), (**B**) *sek-1* (ag1), (**C**) *pmk-1* (km25), (**D**) *skn-1* (zu67), (**E**) *daf-16* (mgDf50), (**F**) *daf-2* (e1368), and (**G**) *dbl-1* (nk3) V. The heat-killed bacterium-treated effects were statistically compared using a log-rank test (N.S. for *P* > 0.05 and \*\*\**P <* 0.0001). (**H**) qRT-PCR Expression of longevity-promoting genes in HK MKAK9-treated worms compared to HK OP50-fed worms. (**I**) A schematic representation showing that HK MKAK9 downregulates insulin-like signaling and upregulates the p38 MAPK pathway to extend longevity in worms. The heat-killed bacterium-treated effects were statistically compared using the Student’s t-test (\**P <* 0.05 and \*\**P <* 0.01). The error bars represent the mean ± SEM.

Subsequent qRT-PCR analysis confirmed the modulation of gene expression involved in insulin-like signaling and p38 MAPK pathway in HK MKAK9-induced longevity. Notably, *daf-2* expression decreased significantly in HK MKAK9-treated worms compared to HK OP50-treated worms (\**P <* 0.05 for *daf-2*) (Fig. 3H). Conversely, the expression of genes involved in the p38 MAPK cascade (*nsy-1*, *sek-1*, and *pmk-1*) increased significantly in HK MKAK9-treated worms (\*\**P <* 0.01 for *pmk-1* and *nsy-1*, \**P <* 0.05 for *sek-1*) (Fig. 3H). Moreover, the expression of the TGF-β homolog *dbl-1* was upregulated approximately two-fold in HK MKAK9-treated worms (\**P <* 0.05 for *dbl-1*) (Fig. 3H). However, longevity assays in *dbl-1* mutants did not support a significant role for *dbl-1* in HK MKAK9-induced longevity.

Furthermore, we confirmed the activation of downstream transcription factors of insulin-like signaling and p38 MAPK cascade, *daf-16* and *skn-1*, by HK MKAK9 treatment. The expression of both *daf-16* and *skn-1* genes significantly increased by approximately two-fold in HK MKAK9-treated worms compared to HK OP50-treated worms (\*\**P <* 0.01 for *daf- 16* and \**P <* 0.05 for *skn-1*) (Fig. 3H). These findings collectively suggest that HK MKAK9 modulates insulin-like signaling and the p38 MAPK pathway to extend longevity in worms (schematically represented in Fig. 3I).

### 3.7 Treatment of HK MKAK9 improves proteostasis by upregulating protein ubiquitination and autophagy receptor expression through insulin-like signaling to extend longevity in worms

We employed mRNA sequencing on young adult wild-type worms following a 24 h treatment with HK MKAK9 and HK OP50 to unravel the downstream pathways of insulin-like signaling and the p38 MAPK pathway modulated by HK MKAK9 in worms. Differential expression analysis of genes revealed a significant transcriptional profile alteration in HK MKAK9-treated worms compared to those treated with HK OP50 (Supplementary file 1, Fig. S5). Subsequently, we generated a heatmap illustrating the differential mRNA expression patterns through hierarchical clustering analyses (Supplementary file 1, Fig. S6). After screening for genes with FDR-adjusted P value < 0.05 and fold change across the mean value of HK MKAK9-treated worms versus HK OP50-treated worms >1, we compiled a list of 229 upregulated and 26 downregulated genes (Supplementary file 2), potentially crucial for HK MKAK9-induced longevity in *C. elegans*.

Protein ubiquitination is a critical regulatory mechanism in proteostasis, involving the tagging of proteins for degradation. This process becomes increasingly important as organisms age and accumulate non-functional proteins. The ubiquitination process is mediated by Cullin-based E3 ubiquitin ligase complexes, prominently including the SCF (Skp, Cullin, F-box containing) complex [37]. These complexes play a pivotal role in recognizing and targeting proteins for ubiquitination and subsequent degradation. At the core of the SCF complex’s functionality are the Skp-1 protein, serving as an adaptor, and various F-box proteins, which are responsible for substrate recognition [37]. Our gene ontology (GO) enrichment analysis revealed upregulation of genes involved in SCF-dependent ubiquitin-mediated protein catabolism, protein ubiquitination, as well as those encoding components of the Cul2-RING ubiquitin ligase complex in HK MKAK9-treated worms compared to HK OP50-treated worms (Fig. 4A, B, and C, Supplementary file 3). Additionally, protein domain enrichment analyses using the DAVID tool by querying the Interpro database identified overexpression of several domain classes related to the SCF complex (SKP1 component (IPR001232), SKP1 component POZ (IPR016073), E3 ubiquitin ligase, SCF complex (IPR016897), SKP1 component dimerization (IPR016072), F-box domain cyclin like (IPR001810), and F-box associated domain type 2 (IPR012885)), further corroborating our findings (Fig. 4D, Supplementary file 3). KEGG analyses highlighted enrichment of critical pathways including ubiquitin-mediated proteolysis, protein processing in the endoplasmic reticulum, and pathways associated with longevity regulation (Fig. 4D, Supplementary file 3). Furthermore, qRT-PCR validation confirmed the upregulated expression of Skp-1-related genes of the SCF complex (*skr*-7, *skr*- 8, *skr*-9, *skr*-14, *skr*-10, *skr*-12, and *skr*-13) involved in protein ubiquitination (\*\**P <* 0.01 for *skr*-8 and *skr*-9, \**P <* 0.05 for *skr*-7, *skr*-14, *skr*-10, *skr*-12, and *skr*-13) (Fig. 5A).

**Figure 4.**
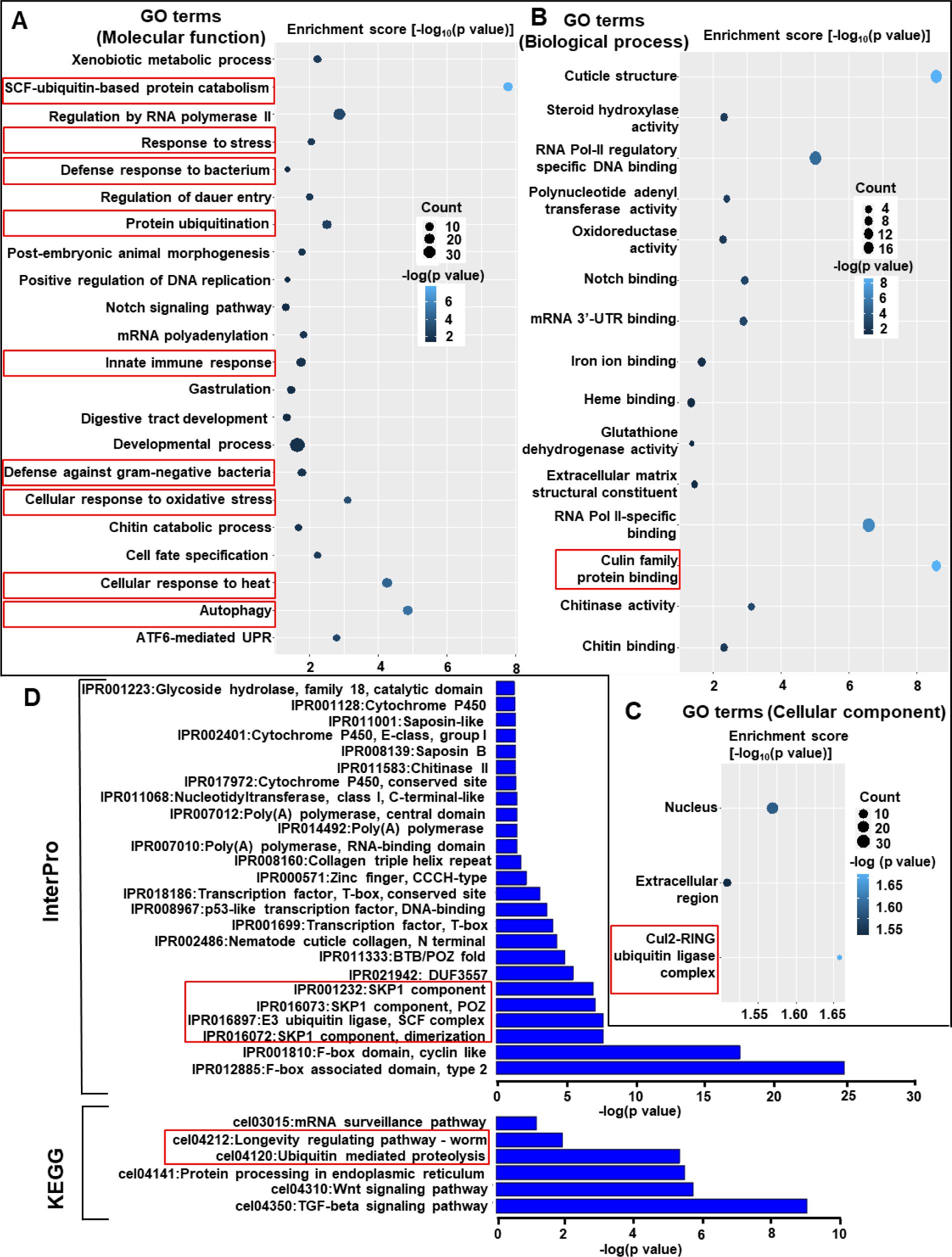
(**A-C**) Gene ontology (GO) analysis of the upregulated genes involved in molecular function, biological process, and cellular components after treatment of HK MKAK9-treated worms compared to HK OP50-treated worms. (**D**) Protein domain enrichment analyses using the DAVID tool by querying the Interpro database and KEGG analysis showing critical pathways.

**Figure 5.**
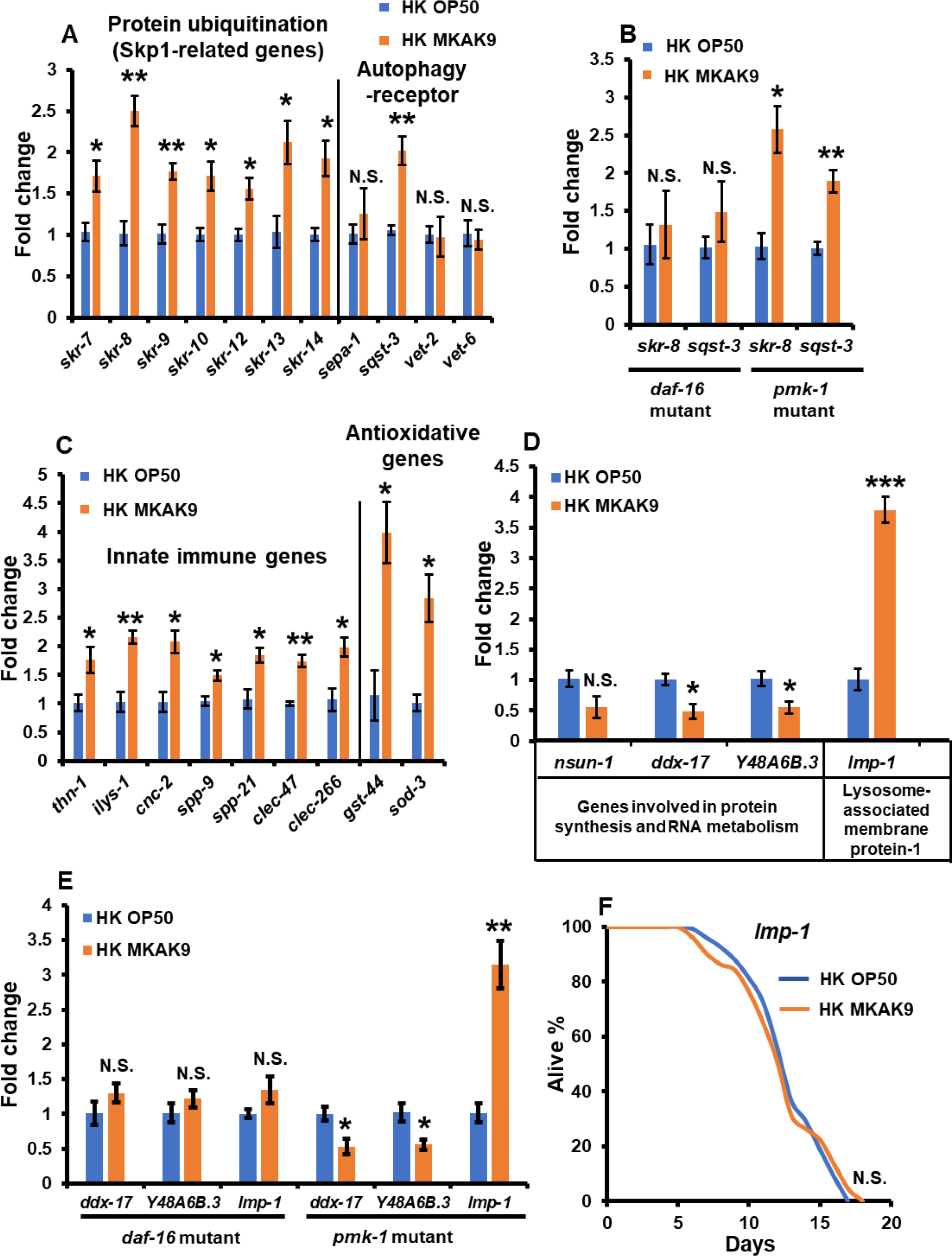
Treatment of HK MKAK9 regulated insulin-like signaling pathway to promote protein ubiquitination and autophagy-lysosome pathway to promote longevity in worms. (**A**) qRT-PCR expression of genes involved in protein ubiquitination (Skp-1-related genes, and autophagy-receptor in wild-type N2 worms. (**B**) qRT-PCR of protein ubiquitination representative gene (*skr-8*) and autophagy gene (*sqst-3*) in *daf-16* (i.e., the downstream target of *daf-2*/insulin-like signaling) and *pmk-1* (i.e., a gene involved in p38 MAPK cascade) mutants (**C**) qRT-PCR expression of genes involved in innate immunity and antioxidation. (**D**) qRT-PCR expression confirmed proteomic analysis results of the genes involved in protein synthesis, RNA metabolism, and the lysosome-associated membrane protein gene lmp-1 (*lmp- 1*). (**E**) qRT-PCR of protein synthesis, RNA metabolism, and the lysosomal protein gene (*lmp- 1*) in *daf-16* and *pmk-1* mutants. (**F**) The lifespan assay on loss-of-function *lmp-1* mutants confirms their role in HK MKAK9-induced longevity in worms. The lifespan assays were statistically compared using a log-rank test (N.S. for *P* > 0.05 and \*\*\**P <* 0.0001). The heat- killed bacterium-treated effects were statistically compared using Student’s t-test (N.S. for *P* > 0.05, \**P <* 0.05, and \*\**P <* 0.01). The error bars represent the mean ± SEM.

Additionally, our gene ontology (GO) enrichment analysis and qPCR results showed upregulation of the *sqst-3* gene encoding an autophagy receptor (\*\**P <* 0.01 for *sqst-3* and N.S. P > 0.05 for *sepa-1*, *vet-2,* and *vet-6*), suggesting the involvement of autophagy in HK MKAK9-induced longevity (Fig. 5A, Supplementary file 3).

Further investigation using qRT-PCR on *daf-16* (downstream target of *daf-2*/insulin-like signaling) and *pmk-1* (a gene involved in the p38 MAPK cascade) mutants revealed that HK MKAK9 regulates protein ubiquitination and autophagy. The results confirmed that HK MKAK9 treatment to *daf-16* mutant did not change the expression of *skr-8* (a protein ubiquitination representative gene) as well as *sqst-3* (an autophagy-receptor) (Fig. 5B) (N.S. *P* > 0.05 for *skr-8* and *sqst-3* in the *daf-16* mutant). In contrast, there was a significant increase in the expression of *skr-8* and *sqst-3* in *pmk-1* mutants (\**P <* 0.05 for *skr-8* and \*\**P <* 0.01 for *sqst-3* in *pmk-1* mutant) (Fig. 5B). Thus, it was confirmed that HK MKAK9 regulated the insulin-like signaling pathway in a DAF-16-dependent manner, not the p38 MAPK pathway, to promote protein ubiquitination and autophagy.

### 3.8 Administration of HK MKAK9 enhances immune responses in worms

GO analysis highlighted upregulation of genes associated with innate immune responses, defense responses to pathogens, as well as cellular responses to oxidative stress and heat (Fig. 4A). Protein domain enrichment analysis indicated increased domain proteins related to immunity, including F-box protein, saponins, and cytochrome P450 in HK MKAK9-treated worms (Fig. 4D). qRT-PCR validation confirmed the upregulated expression of genes involved in innate immunity (*thn*-1, *ilys*-1, *cnc-2*, *spp-9*, *spp*-21, *clec*-47, and *clec*-266) and antioxidative genes (*gst*-44 and *sod*-3) (\*\**P <* 0.01 for *ilys*-1 and *clec*-47, \**P <* 0.05 for *thn*-1, *cnc-2*, *spp-9*, *spp*-21, *clec*-266, *gst-44*, and *sod-3*) (Fig. 5C). These findings shed light on the multifaceted molecular mechanisms underlying HK MKAK9-induced longevity and its potential implications in immunity and antioxidative machinery in *C. elegans*.

### 3.9 Treatment of HK MKAK9 enhanced lysosome-Associated Membrane Protein (LAMP) family protein lmp-1 expression through insulin-like signaling to promote longevity in worms

In our investigation of the effects of HK MKAK9 on proteome dynamics, proteomics analysis was conducted on young adult wild-type N2 worms post 24 h exposure to HK MKAK9 and HK OP50 at 20 °C. This analysis revealed the presence of 2768 proteins across both conditions. Focusing on proteins with significant changes in abundance (based on the median log2-fold change of HK MKAK9/HK OP50), we identified 26 proteins with significant differential expression in the HK MKAK9-treated group (Supplementary file 4). Notably, this group exhibited a marked increase in the abundance of lysosome-associated membrane protein (LAMP) family member lmp-1, a protein synonymous with mammalian lysosomal markers (Supplementary file 4). Subsequent qRT-PCR analysis corroborated the upregulation of *lmp-1* gene expression (\*\*\**P <* 0.001 for *lmp-1*) (Fig. 5D).

Furthermore, the proteomics analysis revealed a decrease in proteins associated with the translational machinery and RNA metabolism in HK MKAK9-treated worms compared to HK OP50-tretaed. This includes proteins such as 26S rRNA {cytosine-C(5)}- methyltransferase (nsun-1), RNA helicase (ddx-17), and a putative H/ACA ribonucleoprotein complex subunit 2-like protein (Y48A6B.3) (Fig. 5D, Supplementary file 4). qRT-PCR analysis confirmed significant downregulation of genes *Y48A6B.3* and *ddx-17* in HK MKAK9- treated worms, whereas *nsun-1* levels did not show substantial alteration (N.S. *P* > 0.05 for *nsun-1*, \**P <* 0.05 for *Y48A6B.3* and *ddx-17*) (Fig. 5D).

To further delineate the regulatory pathways involved, qRT-PCR was performed on *daf-16* and *pmk-1* mutants subjected to HK MKAK9 treatment, aiming to verify its role on the expression of genes linked to protein lmp-1 expression, translational machinery, and RNA metabolism. The analysis revealed no significant changes in the expression of *lmp-1*, *Y48A6B.3*, and *ddx-17* in *daf-16* mutant, whereas *pmk-1* mutant exhibited increased expression of these genes (N.S. *P* > 0.05 for *lmp-1*, *ddx-17*, and *Y48A6B.3* in *daf-16* mutant, \**P <* 0.05 for *ddx-17*, *Y48A6B.3* in *pmk-1* mutant, and \*\**P <* 0.01 for *lmp-1* in *pmk-1* mutant) (Fig. 5E). These findings highlight the pivotal role of the insulin-like signaling pathway, particularly through DAF-16, in regulating the expression of proteins involved in lysosome integrity, protein synthesis, and RNA metabolism in HK MKAK9-treated worms.

Additionally, lifespan assays were conducted on *lmp-1* loss-of-function mutant fed with HK MKAK9 to confirm its role in HK MKAK9-induced longevity. The result indicated that HK MKAK9 treatment did not extend longevity in the *lmp-1* mutant (N.S. *P* > 0.05, log-rank test) (Fig. 5F). In sum, our findings substantiate the hypothesis that HK MKAK9 treatment regulates insulin-like signaling pathways that increase the expression of polyubiquitinated proteins, tagging damaged or aggregated proteins for degradation by activating the autophagy-lysosome pathway to promote longevity in worms.

### 3.10 mir-243 partially lengthens the worm’s lifespan by regulating insulin-like signaling and its downstream genes involved in protein ubiquitination, autophagy receptor Sqst-3, and lysosomal LAMP family protein lmp-1

We next asked whether observed differential gene expression patterns and altered proteomic profiles might be associated with the changes in the microRNA (miRNA) profiles. Through microRNA sequencing, we identified a specific pattern of miRNA expression in HK MKAK9- treated worms: three miRNAs (mir-243, mir-253, and mir-78) were upregulated, while mir- 1818 was downregulated (Supplementary file 5). Subsequent lifespan assays on worms with mutations in these differentially expressed miRNAs revealed that HK MKAK9 treatment significantly enhanced the lifespan in mir-78, mir-253, and mir-1818 mutants (Figs. 6B, C, and D) (\*\*\**P <* 0.0001 for mir-78, mir-253, and mir-1818, log-rank test). Interestingly, while mir- 1818 mutants showed a higher mortality rate than the other miRNA mutants, their lifespan still increased in HK MKAK9-treated worms (Fig. 6D) (\*\*\**P <* 0.0001, log-rank test). However, mir-243 mutant stands out as a distinctive loss-of-function miRNA mutant, exhibiting a noteworthy alteration in the response to HK MKAK9 treatment. Specifically, we found that HK MKAK9-induced longevity was partially declined in the mir-243 mutant and increased only 12.16% in HK MKAK9-treated worms compared to the HK OP50-treated worms (\*\**P < 0.001*, log-rank test) (Fig. 6A). This observation underlines the partial involvement of mir-243 in mediating the longevity-promoting effects of HK MKAK9.

**Figure 6.**
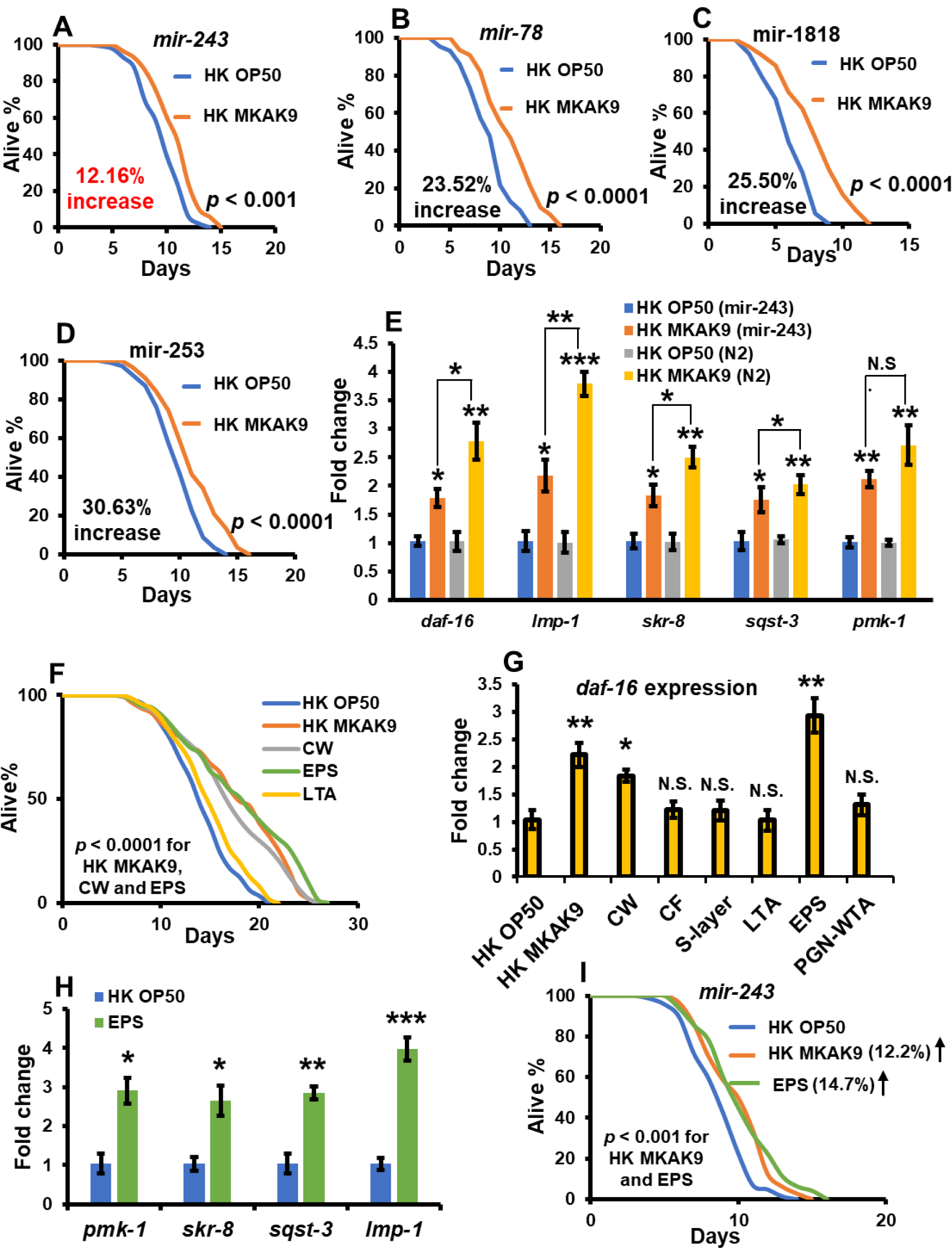
**Treatment of HK MKAK9 and its exopolysaccharide (EPS) partially extend longevity by improving the expression of mir-243 in worms. (A-D**) Lifespan assays on differentially expressed loss-of-function miRNA mutants. (**E**) qRT-PCR expression of genes involved in insulin-like signaling (*daf-16*), protein ubiquitination representative gene (*skr-8*), autophagy-receptor (*sqst-3*), lysosomal protein lmp-1 (*lmp-1*), and the p38 MAPK pathway gene (*pmk-1*) in mir-243 mutant compared to their expression in wild-type N2 worms (Data from Figs. 3H, 5A and D). (**F**). The lifespan assay showing the effect of cellular components and derived EPS component of HK MKAK9 in improving longevity. (**G**) qRT-PCR expression of *daf-16* gene in worms treated with cellular components of HK MKAK9 and compared with *daf-16* expression in HK OP50-treated worms. (**H**) qRT-PCR expression showing the effect of EPS in regulating expression of genes involved in p38 MAPK pathway (*pmk-1*), protein ubiquitination (*skr-8*), autophagy-receptor (*sqst-3*), and lysosomal protein lmp-1 (*lmp-1*). (**I**) EPS of HK MKAK9 partially decreased longevity in the mir-243 mutant. The lifespan assays were statistically compared using a log-rank test (N.S. for *P* > 0.05, (\*\**P <* 0.001, and \*\*\**P <* 0.0001). The heat-killed bacterial effects were significantly compared using Student’s t-test (N.S. for *P* > 0.05, \**P <* 0.05, \*\**P <* 0.01, and \*\*\**P <* 0.001). The error bars represent the mean ± SEM.

To further dissect mir-243’s involvement, we conducted qRT-PCR analysis on mir-243 mutants to elucidate its role in modulating the insulin-signaling pathway and p38 MAPK signaling, alongside its impact on genes associated with protein ubiquitination (*skr-8*), the autophagy receptor (*sqst-3*), and the lysosomal LAMP protein lmp-1 (*lmp-1*) in worms fed with HK MKAK9 compared to those fed with HK OP50. Additionally, we compared the gene expression profiles between the mir-243 mutant and wild-type N2 worms treated with HK MKAK9 and HK OP50 (Data from Figs. 3H, 5A, and D). Our analysis revealed a significant reduction in the expression of *daf-16*, a downstream target of the DAF-2/insulin-like signaling pathway, in HK MKAK9-treated mir-243 mutants compared to *daf-16* expression in wild-type N2 worms (\**P <* 0.05 for *daf-16* in mir-243 mutant and \*\**P <* 0.01 for *daf-16* in N2 worms) (Fig. 6E). Additionally, we observed a notable decrease in the upregulation of representative genes involved in protein ubiquitination (*skr-8*), the autophagy receptor sqst-3 (*sqst-3*), and the lysosomal protein lmp-1 (*lmp-1*) in HK MKAK9-treated mir-243 mutants compared to their expression in HK MKAK9-treated N2 worms (\**P <* 0.05 for *lmp-1* in mir-243 mutant and \*\**P <* 0.01 for *lmp-1* in N2 worms, \**P <* 0.05 for *skr-8* in mir-243 mutants and \*\**P <* 0.01 for *skr- 8* in N2 worms, and \**P <* 0.05 for *sqst-3* in mir-243 mutants and \*\**P <* 0.01 for *sqst-3* in N2 worms) (Fig. 6E). Furthermore, there was no significant change in the expression of *pmk-1*, a gene associated with the p38 MAPK pathway, between mir-243 mutants and N2 worms (\*\**P <* 0.01 for *pmk-1* in both mir-243 mutants and N2 worms) (Fig. 6E). Taken together, our findings from the lifespan screening of differentially regulated miRNA mutants and qRT-PCR analysis indicate that HK MKAK9 upregulates the expression of mir-243, which in turn partially regulates the insulin-like signaling pathway, possibly directly or indirectly modulating protein ubiquitination and their degradation by activating the autophagy-lysosomal pathway to enhance longevity in worms.

### 3.11 Exopolysaccharide (EPS) from HK MKAK9 increases longevity by regulating insulin-like signaling, mir-243 expression, and its downstream genes involved in protein ubiquitination, autophagy receptor (Sqst-3), and lysosomal LAMP family protein lmp-1

We investigated the role of specific cellular components of HK MKAK9 on the longevity of *C. elegans*. Initially, we fractionated various cellular components, including the cell wall, peptidoglycan-wall teichoic acid (PGN-WTA), lipoteichoic acid (LTA), S-layer protein, and extracellular polysaccharides (EPS). We performed lifespan assays with synchronized adult-stage wild-type N2 worms treated with different fractions of HK MKAK9. Specifically, the cell wall and its associated EPS component of HK MKAK9 significantly increased worm longevity by 20.95% and 28%, respectively (\*\*\**P <* 0.0001 for cell wall and EPS, log-rank test) (Fig. 6F). Conversely, treatment with LTA derived from HK MKAK9 did not show a significant effect on worm longevity (*P* > 0.05 for LTA, log-rank test) (Fig. 6F). Additionally, adult wild-type N2 worms cultured on S-layer and PGN-WTA exhibited developmental arrest (Supplementary file 1: Fig. S7). Subsequently, we treated synchronized adult wild-type N2 worms with specific cellular components of HK MKAK9 for 24 h and analyzed the expression of *daf-16* through qRT-PCR. Remarkably, only the cell wall and EPS components of HK MKAK9 induced the upregulation of *daf-16* expression in N2 worms (\**P <* 0.05 for cell wall, \*\**P <* 0.01 for EPS) (Fig. 6G), confirming their role in modulating the insulin-like signaling pathway in a DAF-16-dependent manner. Further analysis revealed that the EPS component of HK MKAK9 also upregulated the expression of *pmk-1* (a gene involved in the p38 MAPK pathway), *skr-8* (representative of protein ubiquitination), *sqst-3* (autophagy receptor), and lysosomal LAMP protein *lmp-1* (\**P <* 0.05 for *pmk-1* and *skr-8*, \*\**P <* 0.01 for *sqst-3*, and \*\*\**P <* 0.001 for *lmp-1*) (Fig. 6H). Lastly, we conducted a lifespan assay using EPS from HK MKAK9 in the mir-243 mutant. Interestingly, EPS only partially increased longevity in the mir-243 mutant, suggesting the involvement of mir-243 in extending the lifespan of wild-type N2 worms (Fig. 6I) (\*\**P <* 0.001, log-rank test). In conclusion, EPS derived from HK MKAK9 promotes longevity by regulating insulin-like signaling (DAF-16-dependent manner) and mir- 243 expression, which further enhances protein ubiquitination and its degradation by activating the autophagy-lysosomal pathway.

## 4. Discussion

The United Nations’ projection for 2050 highlights a significant demographic shift, with one in six individuals expected to be over 65, compared to one in eleven in 2019 [2]. This underscores the pressing need for innovative strategies to address age-related health challenges, including those related to immune responses. Consequently, researchers around the globe are intensifying their efforts on understanding and advancing one key aspect of aging hallmark ‘gut microbiota’ and are exploring microbiota-based therapies to combat disorders linked with aging and age-associated decline in immune function. While much attention has been directed towards live probiotic bacteria and their metabolites [38], the therapeutic potential of heat-killed probiotic bacteria and their structural components remains largely unexplored. In this study, the probiotic bacteria *Levilactobacillus brevis* strain MKAK9 (abbreviated as MKAK9) isolated from mixed pickle was evaluated for its longevity and health-promoting potential using *Caenorhabditis elegans*, an established invertebrate model for studying longevity and conserved aging mechanisms.

Our research revealed that the heat-killed probiotic strain MKAK9 and live probiotic bacteria could lengthen worms’ longevity. Previous studies showed that aging is associated with a decline in age-associated biomarkers in worms, including pharyngeal pumping rate, locomotory activity, and aging pigment lipofuscin accumulation, as reviewed by Son et al. [39]. According to our findings, HK MKAK9 therapy enhanced the age-associated biomarkers in worms. Remarkably, the worms’ survival was prolonged by the administration of HK MKAK9 by reducing pathogens’ colonization and shielded them against *S. aureus* and *E. coli*. Moreover, the HK MRKAK9 therapy augmented the worms’ resistance to heat and oxidative stress. These findings collectively indicate that treatment with HK MKAK9 not only enhances aging-associated biomarkers and extends worm lifespan but also fortifies them against both pathogenic infections (*S. aureus* and *E. coli*) and abiotic (oxidative and thermal) stressors.

Aging is a multifactorial process and involves a complex interplay of various molecular pathways, with prominent roles assigned to key mechanisms such as the insulin-like signaling (IlS) (DAF-2/DAF-16 pathway), the p38 mitogen-activated protein kinase (p38 MAPK) pathway, and the transforming growth factor-β (TGF-β) signaling in worms [22, 40]. The IlS pathway, represented by DAF-2/DAF-16, stands as a cornerstone in the regulation of longevity and healthy aging in *C. elegans* [40]. DAF-2, acknowledged as a gerontogene, acts as the primary receptor for insulin/IGF-1, exerting its influence upstream of the IlS pathway. Reduction in IlS signaling, often through decreased *daf-2* expression, prompts activation of the DAF-16 transcription factor. This activation, in turn, orchestrates gene expression patterns conducive to enhanced longevity and bolstered resistance to stressors [41]. In a parallel vein, the p38 MAPK cascade, typically associated with responses to pathogenic infection and inflammation in humans, exhibits significant relevance in *C. elegans* [22, 40]. Triggered by upstream kinases like NSY-1, SEK-1, and PMK-1, the p38 MAPK pathway encodes a repertoire of antimicrobial proteins and peptides upon encountering bacterial infections [22, 40]. Moreover, PMK-1 activation may engage SKN-1, a pivotal stress-responsive transcription factor that oversees the production of antioxidant proteins via phase-2 detoxification enzymes. By stimulating these immune effectors and antioxidative pathways, activation of the p38 MAPK cascade potentially contributes to lifespan extension [27]. Lastly, the TGF-β pathway emerges as a pivotal player in regulating various aspects of organismal physiology, including longevity, defense against pathogenic infections, dauer formation, and axonal navigation. Its involvement underscores the intricate nexus between developmental processes, immune function, and the aging trajectory in *C. elegans* [42].

In our study, we found that administering the strain HK MKAK9 did not increase longevity in loss-of-function mutants of the p38 MAPK pathway, its downstream gene *skn-1*, and the insulin-like signaling pathways (*daf-2* and *daf-16*) in worms. However, HK MKAK9 treatment did extend longevity in TGF-β homolog *dbl-1* mutant worms. Our qRT-PCR results supported these findings, showing that HK MKAK9 treatment modulated insulin-like signaling (downregulating *daf-2* and upregulating *daf-16*) and increased the expression of genes associated with the p38 MAPK pathway. This concluded that HK MKAK9’s mechanism for extending longevity in worms involves downregulating the insulin-like signaling pathway and upregulating the p38 MAPK pathway.

Our prior research has highlighted the probiotic *Lactobacillus plantarum* JBC5’s ability to activate the p38 MAPK pathway, thereby enhancing worm survival against pathogenic infections by upregulating genes associated with innate immunity [27]. In this study, we further investigated the effects of treating worms with HK MKAK9, elucidating its activation of the p38 MAPK signaling pathway and downstream gene cascades. Gene ontology (GO) enrichment analysis revealed that HK MKAK9 treatment led to the upregulation of genes involved in innate immune responses and defense against pathogens compared to HK OP50- treated worms. qRT-PCR results corroborated these findings, showing increased expression of genes related to innate immunity (*thn*-1, *ilys*-1, *cnc-2*, *spp-9*, *spp*-21, *clec*-47, and *clec*-266) upon HK MKAK9 treatment. These results further support the role of HK MKAK9 in activating p38 MAPK and protecting worms against pathogens like *S. aureus* and *E. coli*. Additionally, protein domain enrichment analysis using the DAVID tool revealed overexpression of several classes, including saponins and cytochrome P450, commonly associated with innate immune responses in worms [22, 40]. This finding aligns with previous research indicating that treatment with probiotic *L. rhamnosus* similarly increases saponins and cytochrome P450 in worms [43]. Next, KEGG pathway analysis demonstrated a 9-fold enrichment for the ‘TGF-β signaling pathway’ and a greater than 5-fold enrichment for the ‘Wnt-signaling pathway’, both encoding molecular effectors crucial for innate immunity [44, 45].

Reactive oxygen species (ROS) levels increase with age and have been linked to over 60 diseases, including gastrointestinal diseases, neurodegenerative conditions, and rheumatoid arthritis [46]. There is compelling evidence suggesting that treatment with HK MKAK9 can decrease ROS levels in worms. Firstly, experiments demonstrate that HK MKAK9 treatment enhances worm survival against oxidative and thermal stress compared to HK OP50-treated worms. Secondly, gene ontology (GO) enrichment analysis indicates the upregulation of genes involved in cellular responses to oxidative stress and heat. Thirdly, experiments using loss-of-function mutants and qRT-PCR confirm that HK MKAK9 treatment activates p38 MAPK, leading to increased expression of downstream transcription factor SKN-1. SKN-1 induction results in the production of antioxidant proteins via phase-2 detoxification enzymes [22]. Consistent with RNA analysis, qRT-PCR results confirm the upregulation of antioxidative genes (*gst-44* and *sod-3*) in HK MKAK9-treated worms compared to HK OP50-treated worms.

GST-44, a glutathione S-transferase omega, is involved in glutathione biosynthesis and detoxifies compounds to reduce ROS accumulation via phase II detoxification [47]. Meanwhile, superoxide dismutase-3 (SOD-3) in the mitochondrial matrix protects worms from oxidative stress during aging [48]. Notably, SOD-3 serves as a transcriptional target for both SKN-1 and DAF-16 due to their binding sites in the promoter region of all SOD genes [48]. Thus, both insulin-like signaling and the p38 MAPK cascade play roles in SOD expression, protecting against oxidative stress and enhancing immunity in worms. However, the interaction between the p38 MAPK cascade and the insulin-like signaling pathway in regulating the expression of antioxidant genes such as *sod-3* in HK MKAK9-treated worms remains unclear. Fourthly, the cellular redox state is measured by SOD activity and the glutathione (GSH) to oxidized glutathione (GSSG) ratio [27]. HK MKAK9 treatment significantly improves SOD activity and the GSH/GSSG ratio while decreasing cytoplasmic ROS compared to HK OP50- treated worms. Fifthly, feeding HK MKAK9 reduces mitochondrial ROS accumulation and increases ATP levels compared to HK OP50-treated worms. In conclusion, these findings support the hypothesis that HK MKAK9 improves cellular and mitochondrial redox states in worms.

Proteostasis is recognized as a significant hallmark of aging, facilitating ubiquitin- mediated proteolysis, which plays a critical role in intracellular protein degradation [49]. Dysfunction in this process has been implicated in various age-associated diseases, including neurodegenerative disorders, sarcopenia, and cardiac dysfunction, as extensively reviewed by Zou et al. [7]. Within the cellular machinery, SKP1, CUL, F-box protein (SCF) complexes represent a subset of SCF E3 ubiquitin ligase complexes responsible for covalently attaching poly-ubiquitin to target proteins, facilitating their efficient elimination via either the autophagy- lysosomal pathway or the ubiquitin-proteasomal pathway [50]. In our study, our mRNA sequencing analysis and qRT-PCR results suggested that HK MKAK9 treatment upregulated the expression of all protein ubiquitination genes (SKR-protein encoding genes) and autophagy-receptor *sqst-3*. Autophagy, a tightly regulated process, plays a pivotal role in degrading improperly synthesized or damaged proteins by enveloping them within double-membrane vesicles (autophagosomes), which subsequently fuse with lysosomes containing hydrolytic enzymes [51, 52]. Autophagy decreases with age due to reduced lysosome-associated membrane protein (LAMP) receptor levels and decreased functional and anatomical features of lysosomes [53, 54]. Our proteomic analysis and qRT-PCR results further indicated that treatment with HK MKAK9 increased the abundance of lysosome-associated membrane protein (LAMP) family protein lmp-1. To validate the significance of lmp-1 in longevity, we conducted a lifespan assay with *lmp-1* mutants. Strikingly, HK MKAK9 treatment failed to enhance longevity in the *lmp-1* mutant, underscoring the pivotal role of lmp-1 in mediating the longevity of HK MKAK9-treated worms. Overall, treatment with HK MKAK9 enhance the expression of genes related to protein ubiquitination, making them more accessible for recognition by autophagy receptors (Sqst-3) located on autophagosomes. These autophagosomes, loaded with cargo, subsequently merge with lysosomes through the LAMP protein lmp-1, leading to the formation of autolysosomes through the autophagy-lysosomal pathway. This process promotes protein homeostasis (proteostasis) and prolong the lifespan of worms.

Furthermore, we performed qRT-PCR in *daf-16* (i.e., a downstream target of DAF- 2/insulin-like signaling) and *pmk-1* (i.e., a gene involved in the p38 MAPK cascade) mutants to understand their role in HK MKAK9-mediated protein ubiquitination and degradation of ubiquitinylated proteins through the autophagy-lysosome pathway in worms. The results showed that treatment of HK MKAK9 to *daf-16* mutant did not increase the expression of genes involved in protein ubiquitination, autophagy receptor Sqst-3, and lysosomal protein lmp-1, while there was significant upregulation in the *pmk-1* mutant. These results suggest that HK MKAK9 regulates the insulin-like signaling pathway for promoting protein ubiquitination and degradation of ubiquitinylated proteins through the autophagy-lysosome pathway in worms. Several lines of evidence support our results: (1) Reduction of insulin-like signaling prevents the upregulation of age-dysregulated deubiquitinating enzymes (DUBs) and promotes the ubiquitination of thousands of proteins, thereby improving protein ubiquitination in worms [50]. (2) Autophagy decreases with age in the pharynx, intestine, neurons, and body wall muscles [55]. Autophagy is crucial for lifespan extension in long-lived insulin-like signaling mutant (*daf-2*) worms and is dependent on the DAF-16 transcription factor [56]. In support, the downregulation of insulin-like signaling or their *daf-2* mutants displayed increased muscle autophagy activity [55]. (3) Li et al. pointed out that reduced insulin-like signaling, including in *daf-2* mutants, reinforces the expression of LMP-1 proteins in a DAF-16-dependent manner, thereby preserving lysosomal integrity and promoting longevity [57]. Our results underline HK MKAK9’s role in downregulating insulin-like signaling pathways (DAF-16-dependent manner) to enhance protein ubiquitination, thereby facilitating the degradation of ubiquitinated proteins through the autophagy-lysosome pathway and extending lifespan in worms.

Our proteomic analysis and qRT-PCR results confirmed a significant decrease in proteins associated with protein synthesis and RNA metabolism, including a putative H/ACA ribonucleoprotein complex subunit 2-like protein (Y48A6B.3) and RNA helicase (DDX-17), following HK MKAK9 treatment. *Y48A6B.3* gene is critical for post-transcriptional modifications, playing a vital role in ribosome biogenesis, protein translation, and pre-mRNA splicing by converting uridine to pseudouridine in target RNAs [58]. DDX-17, an RNA helicase, is essential for translation initiation, ribosome and spliceosome assembly, and RNA structure modification [59, 60]. Collectively, treating HK MKAK9 reduces insulin-like signaling and downregulate mRNA translation or ribosome biogenesis, potentially contributing to the extended longevity of the worms. Supporting evidence indicates that partial inhibition of translation-related processes, such as initiation, elongation, and ribosome biogenesis, through RNA interference (RNAi) enhances longevity and stress resistance in worms [61–69]. Several hypotheses may explain how HK MKAK9 treatment reduces protein synthesis and promotes longevity. First, due to protein folding and translation errors, approximately 30–50% of the nascent polypeptides are co-translationally broken down by ALP or UPS [70, 71]. Thus, reducing global translation might decrease the burden on protein degradation, repair machinery, and maintain protein homeostasis. Secondly, restricting excess biosynthetic and reproductive pathways could help save energy and redirect available resources towards cellular maintenance and repair functions, thereby promoting longevity [63, 72–74].

Epigenetic modifications, such as non-coding RNA (miRNAs), represent another vital hallmark of aging [4]. They contribute to the aging process by modulating gene expression and influencing the cellular mechanisms associated with longevity [3]. MicroRNAs (miRNAs) are key players in a wide array of biological processes including development, aging, cancer, cell- to-cell communication, and physiological responses, as reviewed by Francis et al. [75]. Investigating miRNAs in invertebrate models offers a valuable framework for understanding their roles in more complex vertebrate systems, including humans. Our research utilized microRNA sequencing and lifespan assays on loss-of-function miRNA mutants to uncover that HK MKAK9-induced longevity was notably partially reduced in the mir-243 mutant compared to worms treated with HK OP50. A comparison of qRT-PCR results obtained from mir-243 mutant and wild-type N2 worms confirmed that HK MKAK9 increased mir-243 expression by regulating the insulin-like signaling pathway to partially increase longevity. We next performed qRT-PCR and found that there was a significant reduction in the expression of genes involved in protein ubiquitination, autophagy receptor (*sqst-3*), and lysosomal LAMP protein lmp-1 (l*mp-1*) in HK MKAK9-treated mir-243 mutants compared to their expression in MKAK9-treated N2 worms. Overall, these results confirmed that mir-243 regulates insulin-like signaling to partially increase longevity through modulation of protein ubiquitination and the autophagy-lysosome pathway in worms.

To date, studies focusing on the role of critical postbiotic effectors such as peptidoglycan-wall teichoic acid (PGN-WTA), lipoteichoic acid (LTA), S-layer protein, and exopolysaccharides (EPS) have not been reported about their role in longevity [76]. Using a lifespan assay and qRT-PCR analysis, we discovered that HK MKAK9’s cell wall component and its derived EPS could extend lifespan by attenuating the insulin-signaling pathway in a DAF-16-dependent manner compared to HK OP50-treated worms. Our findings indicate that EPS treatment from HK MKAK9 also increases the expression of genes involved in protein ubiquitination (*skr-8*), autophagy receptor (*sqst-3*), lysosomal protein (*lmp-1*), and the p38 MAPK pathway (*pmk-1*). Furthermore, we observed that the longevity benefits conferred by HK MKAK9 and its EPS was decreased in mir-243 mutants, suggesting a direct or indirect role of mir-243 in promoting partial longevity in worms. Particularly, EPS from *L. brevis* MKAK9, serving as a singular carbon source due to its prebiotic characteristics. *L. brevis* generally produces a water-soluble heteropolysaccharide composed of glucose and sucrose [77–80]. Given its widespread use in fermented foods and the dairy industry, EPS from lactic acid bacteria (LAB) is recognized as safe (GRAS) [77]. LAB-produced EPS is known for its cholesterol-lowering, anti-inflammatory, anti-viral, anti-cancer, and antioxidative benefits [77]. Our primary focus was indeed to contrast the effects of MKAK9’s EPS with those of OP50-treated worms. Therefore, we did not directly compare the obtained results with EPS derived from OP50. Consequently, the distinctions between the EPS from *L. brevis* MKAK9 and OP50 remain to be elucidated. Future investigations will delve into unraveling the structural and functional aspects of EPS from HK MKAK9 and its comparison with EPS derived from OP50 and other *L. brevis* strains. This exploration will contribute to a deeper understanding of the molecular mechanisms underlying longevity and immune response enhancement across various model organisms, both invertebrate and vertebrate.

## 5. Conclusions

Our study demonstrates that the administration of HK MKAK9 and its exopolysaccharides (EPS) enhances longevity and immune responses through a multifaceted mechanism. This process involves a downregulation of insulin-like signaling (DAF-16-dependent manner) and activated downstream pathways responsible for protein ubiquitination and the autophagy-lysosome pathway (Fig. 7). Crucially, ubiquitinylated proteins are identified by the autophagy receptor Sqst-3, engulfed by autophagosomes, and subsequently fused with lysosome (marked by increased lysosome-associated membrane protein lmp-1 expression) to form autolysosomes for degradation of the selected cargo. Notably, both HK MKAK9 and its EPS induced the upregulation of the microRNA (miRNA) mir-243, which plays a role in partially modulating the insulin-like signaling pathway alongside the downstream protein ubiquitination and autophagy-lysosomal pathway (Fig. 7). Furthermore, the activation of the p38 MAPK pathway and its downstream transcription factor SKN-1 by HK MKAK9 and its EPS contributes to enhanced stress resistance, immune responses, and an improvement in the cellular and mitochondrial redox states of the worms. This includes a reduction in mitochondrial ROS, enhancement in mitochondrial transmembrane potential, and an increase in ATP levels (Fig. 7). Collectively, these findings show the great potential of postbiotic HK MKAK9 and its EPS in fostering healthy aging and in the delay of age-associated diseases in mammalian models.

**Figure 7.**
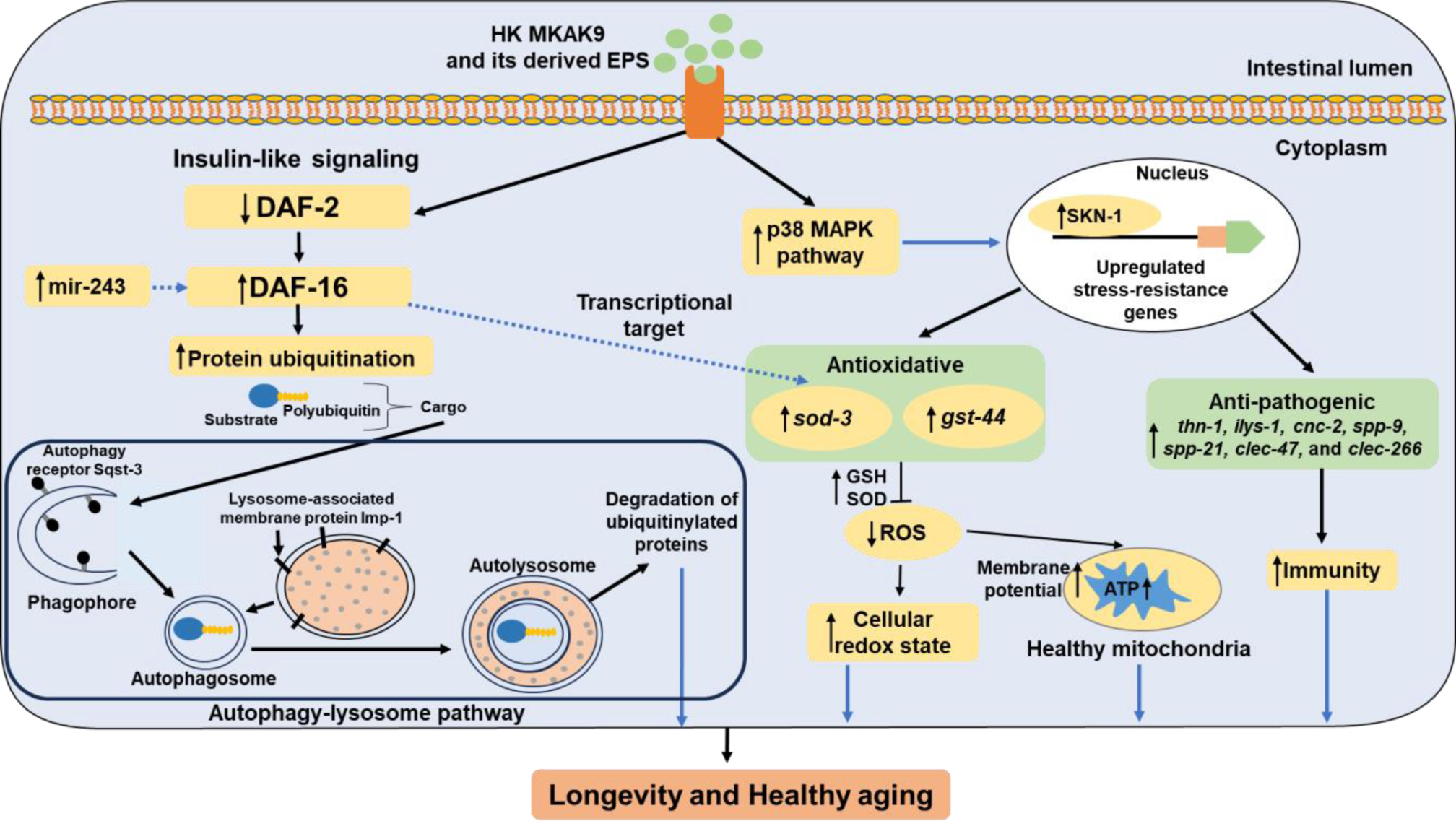
Schematic proposed mechanisms of HK MKAK9 for improving longevity and healthy aging in worms.

## Declarations

### Ethical approval and consent to participate

Not applicable

### Consent for publication

All authors agreed to the submission of this manuscript for publication.

### Availability of data and materials

All data is available in the manuscript, supplementary material, and supplementary files. The 16S rDNA sequence of *Levilactobacillus brevis* MKAK9 has been submitted to the NCBI database (Accession No. OR432299). The raw data generated during mRNA sequencing and small RNA sequencing of treated groups are available at the NCBI Sequence Read Archive (SRA) with accession numbers PRJNA945374 and PRJNA945571. The mass spectrometry proteomics data have been deposited to the ProteomeXchange Consortium via the PRIDE partner repository with the dataset identifier PXD041237.

### Author’s contributions

The authors AK and MRK designed the experiments. AK performed all the experiments. AK, MKS, VP, AB, and SB analyzed the data. The first draft of the manuscript was written by the author AK and further revised by MRK, SWC, AKM, and MCK. All authors have agreed for the publication of a final manuscript.

## Acknowledgement

Not applicable

## Supplementary material legends

**Supplementary file 1** This file contains supporting supplementary figures and tables. **Figure S1.** The evolutionary distance between *Levilactobacillus brevis* strain MKAK9 (Accession no.: OR432299), type strain *Levilactobacillus brevis* strain ATCC 14869 (Accesion no.: NR 044704), and related species of *Lactobacillus* was calculated based on 16S rRNA sequences and neighbour-joining tree was computed. The nodes consist bootstrap value (percentages of 1000 replications) greater than 60%. The parentheses of each strain are represented with the GenBank accession number. An outgroup, *Enterococcus faecium* ATCC 19434 (Accession no.: DQ411813.1) was used to show the evolutionary distances. Kimura 2-parameter method was selected to compute the evolutionary distance in MEGA 7 software, showing the nucleotide substitutions per base. There were a total of 1369 positions in the final dataset. The bar representing 0.01 substitutions per base site. **Figure S2.** The developmental rate (egg-to-egg time) was measured after treatment with HK OP50 or HK MKAK9, as discussed by Soukas et al. [29]. Treatment effects were compared using Student’s t-test (\*\**P* > 0.01). **Figure S3.** The brood size was analyzed after feeding HK OP50 or HK MKAK9. Error bars represent mean ± SEM. Treatment effects were compared using Student’s t-test (N.S. *P* > 0.05). **Figure S4.** (**A** and **B**) Pre-treatment of worms on HK MKAK9 provided resistance against pathogens *S. aureus* MTCC3160 and *E. coli* MTCC 1687. The heat-killed bacterium-treated effects were statistically compared using Student’s t-test (\*\**P <* 0.01, and \*\*\**P <* 0.001). **Figure S5.** Gene transcription profile (volcano plot) of HK MKAK9-treated worms in comparison to HK OP50- treated worms. The red dots showed downregulated genes and green dots represented upregulated genes. The blue dots indicated genes with no significant change in the expression. **Figure S6.** Heat map analysis of RNA sequencing data showing top 20 genes those were upregulated and downregulated in HK MKAK9-treated worms in comparison to HK OP50- treated worms. Each treatment group had three biological replicates. Each group has three single neurons tested. The red colors indicate high expression genes and blue colors denotes the low expression genes, respectively. The log2 value was calculated from mean signals. **Figure S7.** The effect of specific cellular components of HK MKAK9 on development of young adult (day 3) wild-type N2 worms. The graph was plotted as percent developed worms after 2 days (day 5) with different treatment groups by counting the number of developed worms. **Table S1** Closest relative of the strain *Levilactobacillus brevis* strain MKAK9 in the NCBI database. **Table S2** Primer sets designed for qRT-PCR analysis in this study. **Table S3** Survival and adhesion of MKAK9 under *in vitro* simulated gastro-intestinal conditions. **Table S4** Antibiotic sensitivity of the strain MKAK9. **Table S5.** The mean lifespan of C. elegans strains treated with HK MKAK9 or HK OP50 (N.S. for *P* > 0.05 and \*\*\**P* < 0.0001, log-rank test).

**Supplementary file 2.** A list of differentially upregulated and downregulated mRNA genes

**Supplementary file 3.** Gene ontology enrichment analysis and a list of differentially expressed genes belonging to biological processes, molecular functions, and cellular components.

**Supplementary file 4.** A list of differentially regulated proteins in treated groups.

**Supplementary file 5.** A list of differentially expressed microRNA in treated groups.

